# Enhanced forest heterogeneity drives stronger functional than taxonomic shifts in soil nematodes

**DOI:** 10.1101/2025.08.30.673220

**Authors:** Rike Schwarz, Pia M. Bradler, Anne Chao, Marcel Ciobanu, Orsi Decker, Benjamin M. Delory, Peter Dietrich, Andreas Fichtner, Ludwig Lettenmaier, Michael Junginger, Oliver Mitesser, Jörg Müller, Goddert von Oheimb, Kerstin Pierick, Nico Eisenhauer, Simone Cesarz

**Affiliations:** German Centre for Integrative Biodiversity Research (iDiv) Halle-Jena-Leipzig, Leipzig, Germany; Institute of Biology, Leipzig University, Leipzig, Germany; Institute of Ecology, Leuphana University, Germany; Institute of General Ecology and Environmental Protection, TUD Dresden University of Technology, Tharandt, Germany; Institute of Statistics, National Tsing Hua University, Hsin-Chu, Taiwan; Institute of Biological Research, Branch of the National Institute of Research and Development for Biological Sciences, 48 Republicii Street, 400015, Cluj, Napoca, Romania; Bavarian Forest National Park, Grafenau, Germany; Environmental sciences, Copernicus institute of sustainable development, Utrecht University, The Netherlands; Institute of Biology / Geobotany and Botanical Garden, Martin Luther University Halle-Wittenberg, Halle, Germany; Chair of Conservation Biology and Forest Ecology, Biocenter, University of Würzburg, Rauhenebrach, Germany; Silviculture and Forest Ecology of the Temperate Zones, University of Göttingen, Germany

## Abstract

Production forests are often managed primarily for timber production, leading to biotic homogenization and reduced biodiversity. To explore strategies that promote biodiversity while maintaining timber yields, we conducted a large-scale experiment in eight German forests. We manipulated structural β-complexity, i.e., the heterogeneity of structural elements across forest patches, by experimentally introducing variation in canopy gaps and different types of deadwood across 156 plots of 50 × 50 m each, to investigate its effects on forest biodiversity. We analyzed soil nematode communities, which are important bioindicators and contributors to ecosystem processes, by assessing taxonomic and functional diversity across patch (α), site (γ), and between-patch (β) scales using Hill–Chao numbers. Additionally, we tested whether environmental variables explain nematode diversity responses. Our results show that functional diversity is more responsive than taxonomic diversity, with increased β-diversity of common and frequent taxa alongside simultaneous declines in α– and γ-diversity. This pattern suggests a shift toward more specialized nematode communities in response to the intervention. Moreover, we found that site-specific conditions, such as sand content and understory biomass, modulated these effects. Overall, our findings reveal complex, scale-dependent responses of nematode diversity to aboveground forest structural changes, emphasizing the need to consider environmental context in forest biodiversity management. This study represents an important first step toward understanding and enhancing soil biodiversity at management-relevant spatial scales.

## Introduction

Increasing human pressure decreases biodiversity and changes species compositions globally (Díaz et al., 2019; Keck et al., 2025). Land-use intensification is a key driver of biodiversity loss in terrestrial ecosystems (Jaureguiberry et al., 2022; Newbold et al., 2015). Understanding how biodiversity changes is essential for effective conservation and management. However, biodiversity responses to anthropogenic change are scale-dependent, so focusing solely on local (alpha, α) diversity can obscure important patterns occurring at broader spatial scales (Chase et al., 2020, Dornelas et al., 2014). In particular, beta (β) diversity—representing the dissimilarity between communities—provides critical insights, because it reveals how species composition changes across habitats or regions, helping us understand ecosystem connectivity, drivers of biodiversity patterns, and priorities for conservation (Anderson et al., 2011). While α-diversity may remain stable or even increase, β-diversity can decline, indicating biotic homogenization (Chase et al., 2020; Delgado-Baquerizo et al., 2021; Gossner et al., 2016). Because different communities fulfill distinct functions depending on environmental context (Isbell et al., 2011), shifts in β-diversity can have cascading effects on ecosystem multifunctionality (Mori et al., 2018), underscoring the need to consider diversity across spatial scales.

In addition, species’ responses to environmental change vary depending on their abundance, with rare, frequent, and common species often showing distinct and sometimes contrasting patterns (Basile, 2022; Roswell et al., 2021, 2023). Moreover, biodiversity can be described along multiple facets, including taxonomic and functional dimensions (Cadotte et al., 2011; Jarzyna & Jetz, 2016). Taxonomic diversity captures differences in species identities, while functional diversity groups species based on shared traits and reflects the ecological roles they perform (Cadotte et al., 2011). Importantly, the degree of functional dissimilarity among species has been shown to drive net biodiversity effects on ecosystem functioning (Heemsbergen et al., 2004), most likely due to niche complementarity (Wagg et al., 2017). While functional diversity may change more strongly than taxonomic diversity (Mori et al., 2015), it can also change less when many species fall in the same functional group (i.e., are functionally similar) (Eisenhauer, 2023; Jarzyna & Jetz, 2018). This emphasizes the importance of considering multiple facets (Craven et al., 2018).

Land-use intensification in forests, such as through intensive forestry practices in Europe, has often reduced features typical of early successional stages, like canopy gaps and deadwood (Aszalós et al., 2022; Paletto et al., 2014). We define these elements as key components of structural complexity in a forest (Keeton, 2006; Müller et al., 2023). The reduction of them decreases the structural dissimilarity (= heterogeneity) between stands and can therefore lead to landscape and biotic homogenization (Rousseau et al., 2019). According to the habitat heterogeneity hypothesis, a more complex habitat increases species diversity due to more niches (Heidrich et al., 2020; Stein et al., 2014). In forests, both vertical and horizontal heterogeneity have been identified as major drivers of biodiversity (Heidrich et al., 2020). Enhancing habitat heterogeneity, i.e., by increasing structural complexity through the addition of deadwood and the creation of canopy gaps, may therefore help counteract biodiversity loss (Hekkala et al., 2023; Müller et al., 2023). While numerous studies have investigated the effects of deadwood and/or canopy gaps on forest biodiversity at the stand (α) scale (e.g. Dyson et al., 2024; Penone et al., 2019; Rothacher et al., 2023; Wang, 2025), little is known about how these factors influence biodiversity at the between-stand (β) and landscape (gamma, γ) scales. To explore this overlooked aspect, this study focuses on enhancing structural β complexity (ESBC), which refers to the variation in structural features between stands within a landscape. This includes differences in deadwood types, such as snags, stumps, and lying logs, as well as canopy conditions (Müller et al., 2023). Such variation can foster distinct local communities (α-diversity) due to differing environmental conditions, leading to increased β-diversity between stands (Montoya-Sánchez et al., 2023). In turn, higher β-diversity contributes to greater overall landscape-level (γ) diversity. ESBC may thus be an effective way to increase habitat heterogeneity and biodiversity at management-relevant scales in forests (Müller et al., 2023).

Soil is a central component of forest ecosystems, providing habitat for a vast diversity of organisms (Anthony et al., 2023; Baldrian, 2016). Soil biota play a crucial role in ecosystem functioning by mediating key processes such as decomposition, nutrient cycling, and the redistribution of resources (Bardgett & van der Putten, 2014; van der Heijden et al., 2008). Among soil fauna, nematodes represent a key group (Yeates, 2007) being the most abundant metazoans on Earth (Orgiazzi et al., 2016). Nematodes are commonly used as bioindicators (Biryol et al., 2024; Cesarz et al., 2013; Richter et al., 2023), because they are widely distributed across ecosystems, highly diverse, have short generation times, represent different soil food web components and energy channels, respond rapidly to environmental change and enrichment, and can be identified relatively easily based on morphology (Bongers & Ferris, 1999; van den Hoogen et al., 2019). Nematodes are typically grouped into five major trophic groups: bacterial feeders, fungal feeders, plant feeders, omnivores, and predators. These reflect their position in the soil food web (Yeates & Bongers, 1993). In addition, species are distributed along a colonizer-persister gradient (c-p values), reflecting their life-history strategies, with lower values indicating opportunistic, disturbance-tolerant taxa and higher values indicating more sensitive, slow-reproducing taxa (Bongers & Bongers, 1998). Combining trophic group and c-p classification yields functional guilds that provide a nuanced view of soil ecosystem conditions and help to capture functional changes (Cesarz et al., 2015). The abundance, composition, and diversity of nematode communities are strongly influenced by environmental conditions. These include plant diversity (Dietrich et al., 2021; Eisenhauer et al., 2013; Zhang et al., 2015), and soil properties such as water availability (Franco et al., 2019), organic carbon (Shao et al., 2023), pH (van den Hoogen et al., 2019), nitrogen content, and microbial biomass (Mueller et al., 2016).

Forest management practices that increase canopy openness and deadwood abundance may modify soil abiotic conditions, including water content, pH, and carbon and nitrogen content, as well as the availability and diversity of food resources for nematodes by altering plant, bacterial, and fungal communities (Mueller et al., 2016; Perreault et al., 2020; Wang et al., 2022, 2024). Such environmental changes may cascade into nematode community composition and diversity by altering both habitat conditions and bottom-up resource dynamics (Biryol et al., 2024; Thakur et al., 2014; Yeates, 2007).

In this study, we aim to address the research gap regarding how enhancing structural β complexity in forests affects the taxonomic and functional diversity of rare, frequent, and common nematode taxa on the α, β, and γ level (Jost, 2007).

We hypothesized that:

1. Enhancing structural β complexity in forests increases both functional and taxonomic β-diversity by creating more distinct habitat types, which in turn leads to higher γ-diversity.
2. Effect sizes of enhanced structural β complexity on nematode diversity metrics can be explained by variation in environmental variables, such as deadwood quantity, air temperature, soil pH, soil microbial biomass, understory biomass and species richness, soil carbon, nitrogen and C:N ratio, litter depth, and sand content as a proxy for soil water availability.

## Methods

### Study design

This study was embedded within the BETA-FOR experimental framework (Müller et al., 2023), which was established to assess ecological responses to enhancements of structural β complexity across forests in Germany. Eight predominantly beech-dominated deciduous forest sites were studied. These included one site located in the University of Würzburg’s forest (U03), five sites within or near the Bavarian Forest National Park (B04, B05, B06, B07, P08), one site near Saarbrücken (S10), and one in the vicinity of Lübeck (L11; Figure 1A). Each forest site consists of a control district and a treatment district (Figure 1B). A district (10–20 ha) contains nine 50 x 50 m patches (15 in U03). Control and treatment districts of the same forest site were established in close proximity to each other, ensuring similarity in site conditions such as geology, soil, and tree composition. On the patches within the treatment districts, silvicultural interventions were applied to 30% of the stand basal area to create canopy openings and deadwood. As the aim was to enhance structural β complexity (i.e., ESBC = different kinds of intervention within a district), specific treatments were applied to the ESBC patches of the same district (Figure 1B and 1C).

**Fig. 1:**
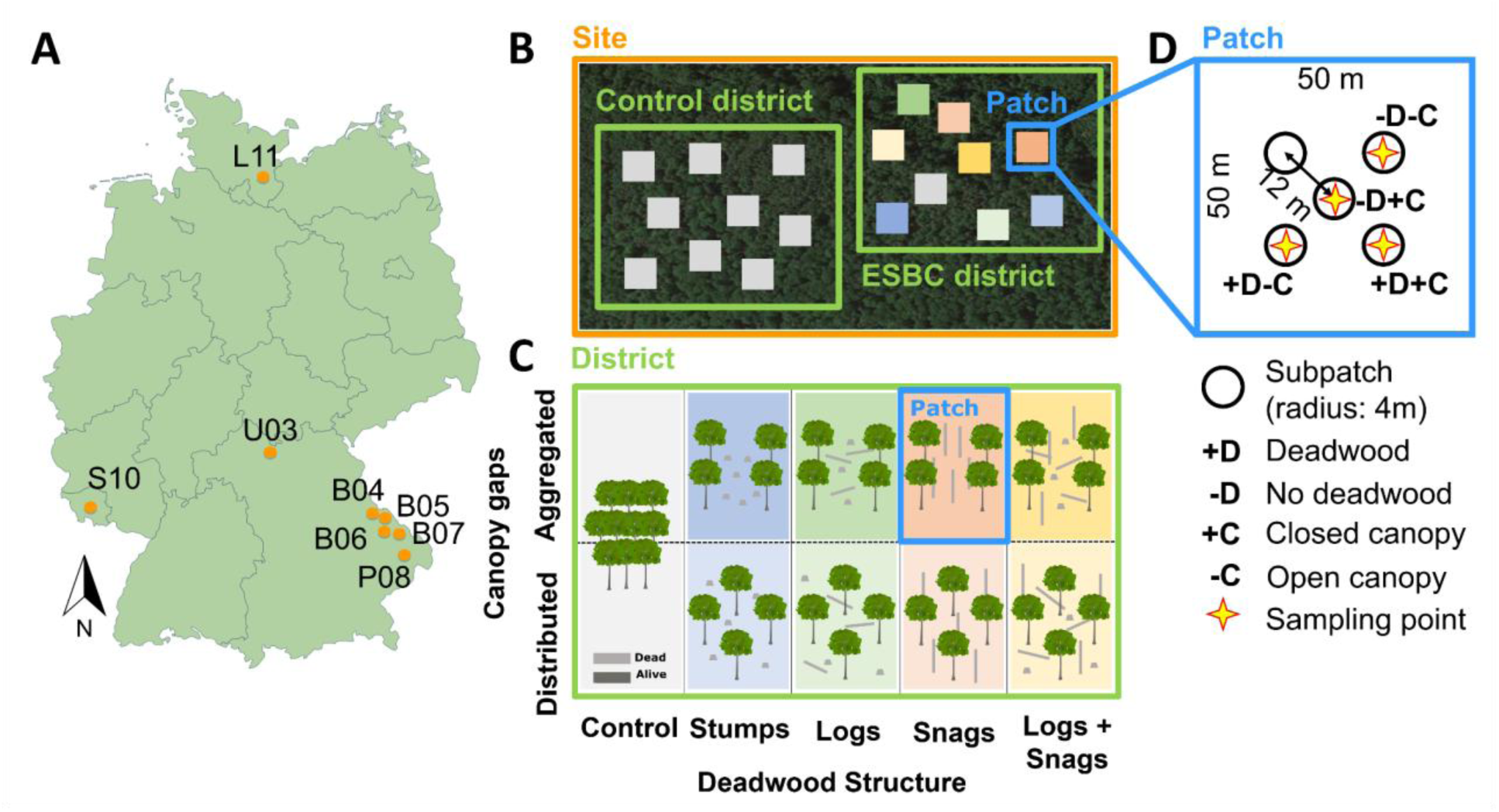
(A) Locations of the study sites across Germany. (B) Schematic representation of a site showing the ESBC (enhancement of structural β complexity) and the control district. (C) Illustration of the individual treatments within an ESBC district along two axes: canopy gaps in aggregated and distributed mode and different deadwood structures. (D) Schematic representation of the spatial sampling within a patch. Within-patch heterogeneity was captured by sampling across combinations of canopy cover (open vs. closed) and deadwood presence. Ideally, one soil sample was taken from each of the following conditions: (1) deadwood under open canopy, (2) no deadwood under open canopy, (3) deadwood under closed canopy, and (4) no deadwood under closed canopy. Sampling began at the patch center, from which three subpatches were selected to best represent all condition combinations, i.e., the heterogeneity of the patch. Figure modified after Schwarz et al., 2025.

**Figure 1:**
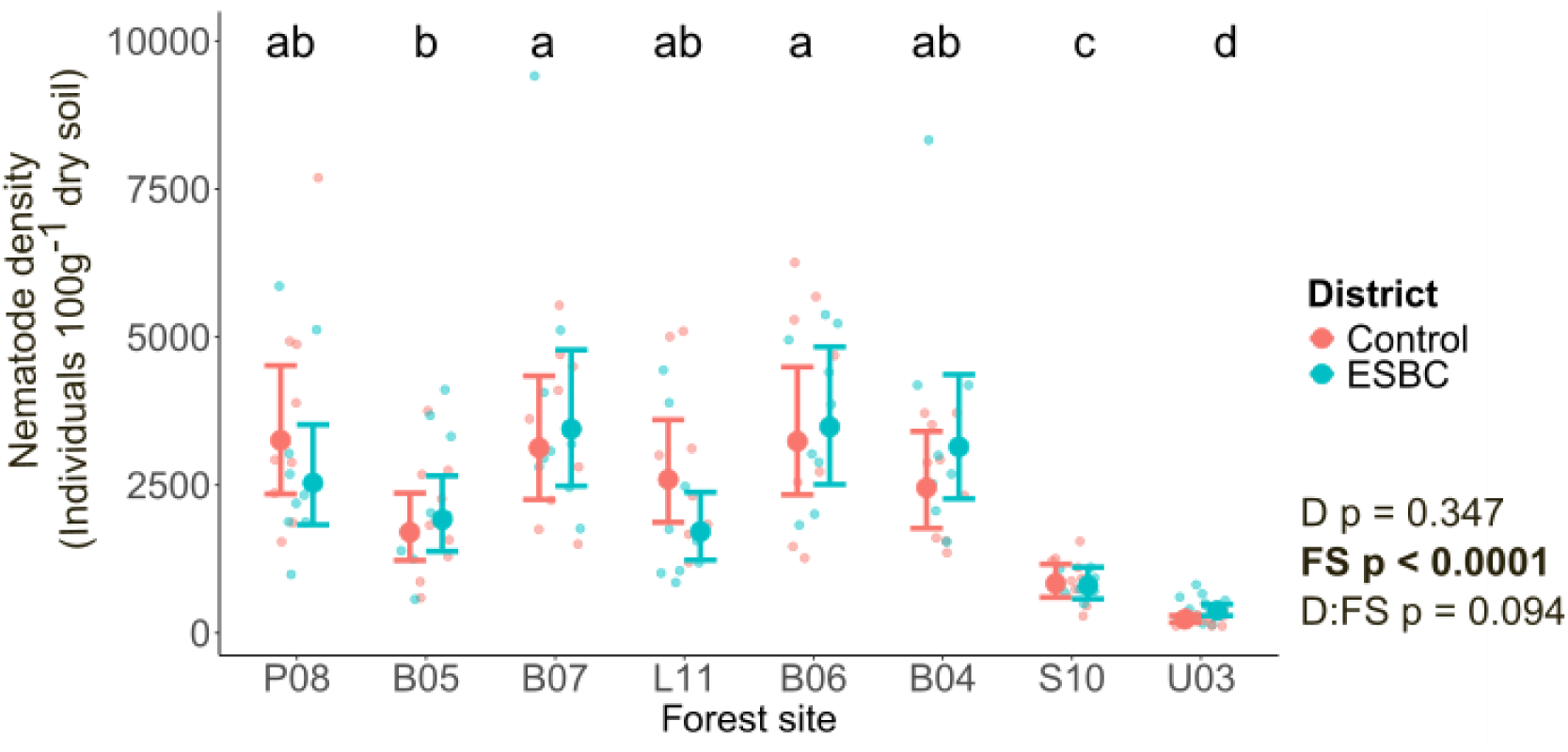
Nematode density (individuals per 100 g dry weight soil) per forest site (FS) and district (D): Control is displayed in red and enhanced structural β complexity (ESBC) in blue. P-values are derived from a linear model with two independent variables with significant values highlighted in bold (p-value < 0.05). Letters represent a significant difference between forest sites derived from a post-hoc test. The model predictions are displayed with error bars representing the 95% confidence intervals and mean values indicated by larger points, while raw data points per patch are shown in the background. Forest sites are ordered by increasing soil pH value.

Therefore, two variants of tree removal were implemented: an aggregated mode, resulting in a single large canopy gap (30 m diameter), and a distributed mode, where tree removals were spread evenly across the patch avoiding large openings. These two modes of canopy openings were crossed with different deadwood treatments: stumps, logs, snags, or logs and snags remaining (Figure 1C). Previous analyses have demonstrated that these ESBC treatments indeed increased forest structural heterogeneity at the landscape scale (Pierick et al., 2025). Forest site U03 included six additional patches accommodating three more deadwood treatments (total tree removal with an excavator, crowns remaining, and habitat trees) to allow investigation of a higher level of spatial heterogeneity, resulting in 15 patches. On average, the interventions led to 45 m³ (± 29 m³) of deadwood per hectare in the ESBC patches. To further increase the heterogeneity of the ESBC district, an untreated patch was added. Control patches also remained untreated, reflecting standard forest management practices or natural development in formerly managed stands within the national park. In total, 156 experimental forest patches (78 ESBC and 78 Control) were investigated in this study.

### Soil sampling and nematode extraction

All patches were sampled in October 2023. After roughly removing the litter, we took four soil cores (5 cm diameter, 10 cm depth) per patch. Sampling points were chosen to represent heterogeneity regarding deadwood and light availability within a patch (Figure 1D). The soil was sieved (2 mm), pooled per patch, and stored at 4°C until further processing. We extracted the nematodes using the Baermann-Funnel method from ∼ 25 g of fresh soil per patch (Cesarz et al. 2019). After 72 h, we emptied the tube containing the nematodes through a 15 µm mesh and fixed the nematodes immediately in 4% formalin. We determined the dry weight of the soil from each sample. Nematode individuals were counted at 50x magnification. A random fraction of the specimens, ideally 100 individuals, were identified at 400× magnification using a Leica DMI 4000B light microscope and following Bongers (1994). Adults and most of the juveniles were identified to genus level; some juveniles were also identified to family level. As some of the nematode samples had lots of debris besides the nematodes, it was not possible to identify 100 specimens in each sample. To account for differences in the number of identified nematode individuals across samples (ranging from 9 to 108), we estimated α-diversity using the iNEXT approach (Chao et al., 2014; Hsieh et al., 2016). iNEXT employs rarefaction and extrapolation to standardize diversity estimates across uneven sample coverages. This approach allows us to include all samples, even those with low numbers of individuals identified, in the diversity comparison while reducing bias due to unequal identification intensity.

Nematode taxa were then classified according to the c–p gradient (Bongers, 1990; Bongers & Bongers, 1998) and the feeding types (Bongers & Bongers, 1998; Okada et al., 2005; Yeates & Bongers, 1993). For the assignment of nematode genera to trophic groups, we primarily referred to Yeates & Bongers (1993). However, since Nemaplex provides the most up-to-date information, all assignments were cross-referenced with the Nemaplex database (http://nemaplex.ucdavis.edu/). Similarly, cp-values were initially assigned based on Bongers (1990) and Bongers and Bongers (1998), with all values verified and updated according to Nemaplex when applicable. Since Nemaplex is a dynamic resource, our specific assignments, as confirmed by Nemaplex on January 17, 2025, are provided in the Supporting Information (Table S1 in Supporting Information). Although Tylenchidae are typically plant feeders, individuals not identified to genus level were classified as fungal feeders in our dataset, as the dominant genus *Filenchus* in our samples is known to feed on fungi (Okada & Kadota, 2003). Nematode average body mass (for calculating functional diversity, see below) was retrieved from Nemaplex on March 24, 2025, and is also provided in the Appendix (Table S1 in Supporting Information).

### Environmental variables

Soil pH was determined by mixing 10 g of air-dried soil with 25 ml of a 0.01 M CaCl₂ solution, followed by shaking and a 1-h settling period. The pH value was then measured using a pH meter (Orion Star A211, Thermo Scientific, MA). For carbon and nitrogen analysis, soil samples were dried at 30°C for 72 h, finely ground, and weighed into tin capsules (20 mg per sample). Total C and N contents were measured through dry combustion using a Vario EL cube IR elemental analyzer. Results were expressed as percentages of the total sample mass, and the C:N ratio was calculated accordingly. Microbial biomass was measured using substrate-induced respiration (Scheu, 1992). An aqueous glucose solution was added to soil samples, and the resulting respiratory response was used to calculate microbial biomass (Beck et al., 1997, p. 199). This reflects the microbial community capable of using glucose as a carbon source. We used soil sand content as an indirect indicator of soil water-holding capacity (Rout & Arulmozhiselvan, 2019). Sand content and litter layer thickness were recorded by an expert during site mapping following the methodology described in an authoritative German reference manual on forest site classification (Forstliche Standortaufnahme, 2016). Deadwood volume was measured across all patches. On the ESBC patches, all experimentally added deadwood (lying, standing, stumps, and habitat trees) was assessed. On the control patches, only naturally occurring standing dead trees and freshly cut stumps resulting from thinning operations were documented. For standing deadwood, a taper rate of 1 mm per 10 cm of height was applied. The volume of tree crowns and lying deadwood was estimated using the 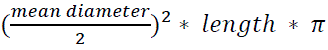. Deadwood with a diameter smaller than 7 cm, including both snags and logs, was excluded, thereby restricting the assessment to coarse woody debris. The term “deadwood” is used throughout this manuscript to represent coarse woody debris. Air temperature at a height of 2 m was recorded on each patch using one EL-USB2 data logger (Lascar Electronics Ltd., UK). Each logger was placed inside a TX COVER shield (Technoline, Germany) and mounted on wooden poles located at the center of the patches. Measurements were taken at 30-minute intervals. Missing data caused by logger malfunctions were estimated using data from loggers within the same district, applying the R package *‘MissForest’* (Stekhoven & Bühlmann, 2012). The resulting temperature records were then aggregated to calculate mean annual air temperature for 2023 for each site. During the 2023 growing season, understory surveys and aboveground biomass sampling were carried out in five subpatches (each with a 4 m radius) within every patch. In the vegetation surveys, all herbaceous and woody plant species up to 1 m in height were recorded. For biomass collection, approximately 0.5 m² (70 × 70 cm) of understory vegetation was harvested from each subpatch at a spot selected to be representative of both species composition and biomass of the patch and subpatch. The collected plant material was oven-dried at 60 °C for 72 h and subsequently weighed. Biomass values were then aggregated for each patch and calculated for an area of 1 m².

### Statistical analysis

We fitted a linear model with forest site, district, and their interaction as explanatory variables to assess whether ESBC affects nematode density per 100 g dry weight and to check if this effect differs between forest sites.

To assess the effect of ESBC on nematode diversity, we analyzed α-(within-patch), γ-(across-district), and β-(between-patch) diversity using Hill numbers, a unified framework of diversity metrics that account for both species richness and species relative abundances. A key advantage of Hill numbers is their ability to provide a consistent interpretation of diversity across different scales (α, β, γ) and orders (q), facilitating direct comparisons and better ecological understanding (Hill, 1973; Jost, 2006, 2007). When q = 0, all species are weighted equally, making the measure equivalent to species richness and, therefore particularly sensitive to rare species. At q = 1, species are weighted according to their relative frequencies, resulting in the exponential of Shannon entropy. This balances the influence of both common and rare species, allowing the diversity to be expressed as the effective number of equally frequent species. When q = 2, the measure gives greater weight to dominant species, corresponding to the inverse of the Simpson index. We applied this approach to both taxonomic and functional diversity. For taxonomic diversity, we used family and genus-level classification of nematodes, from here on referred to as taxa. For functional diversity, we selected three traits: c-p value (colonizer-persister scale), trophic group, and body mass. Trait dissimilarities were calculated using the Gower distance (Legendre & Legendre, 1998; Podani, 1999), a metric that accommodates mixed data types (continuous, ordinal, categorical) (Pavoine et al., 2009). These dissimilarities were then used to generate functional species equivalents, which were treated analogously to taxonomic units in subsequent diversity analyses.

To calculate β-diversity, we used the multiplicative partitioning approach, where β-diversity is defined as the ratio of γ-diversity to α-diversity (β = γ / α) (Jost, 2007; Whittaker, 1972). However, as β-diversity can be biased by unequal sampling completeness across sites, we standardized diversity estimates based on sample coverage—the proportion of the total community that is represented in a sample. When Hill numbers are estimated at a standardized level of sample completeness rather than based on raw sample size, they are referred to as Hill-Chao numbers (Chao et al., 2014; Chao & Jost, 2012). This ensures that diversity comparisons are not biased by differences in sample completeness. In the case of abundance data, extrapolation of diversity estimates is considered robust up to different limits depending on the diversity order: for q = 0, extrapolation is reliable only up to twice the observed sample size (i.e., number of individuals), since extrapolation beyond that becomes highly uncertain due to the difficulty of reliably estimating the many undetected rare species. For q = 1 and q = 2, extrapolation up to complete coverage (100%) is generally accepted (Chao et al., 2023).

As a basis for all diversity analyses, we used raw abundance data rather than extrapolated values (e.g., standardized to total nematode counts or per 100 g dry weight). This was essential, because the method implemented relies on the presence of singletons, which serve as indicators of rare species and are critical for accurate sample completeness estimation. To account for differences in the number of measured assemblages per site (e.g. 9 vs. 15), we used the Jaccard turnover transformation of β-diversity (1–S) (Chao et al., 2019). This transformation expresses β-diversity as the proportion of species turnover (compositional dissimilarity) that is independent of sample size.

For each site, we first calculated the observed sample coverage (Figure S1 A in Supporting Information). Based on these values, we defined two thresholds for extrapolation. The first threshold, Cmax (conservative), corresponds to the maximum level of coverage that could be achieved by doubling the sample size of the forest site with the lowest observed completeness, thereby providing a conservative standard for cross-site comparisons. The second threshold, C25 (inclusive), was defined as the 25th percentile of extrapolated coverage values across all sites, representing a more inclusive standard that allows comparisons at higher coverage levels. In our dataset, Cmax was 0.849 and C25 was 0.974 (Figure S1 B in Supporting Information). For our main analysis, we used Cmax, but all diversity estimates were computed at both levels to assess the robustness of our conclusions.

To evaluate ESBC effects statistically, we calculated site-level differences in response to ESBC for each diversity metric and performed a fixed-effect meta-analysis. To ensure robust and reproducible estimates, we used 300 bootstrap replicates to compute standard errors, from which 95% confidence intervals were derived. This approach allowed us to estimate the overall treatment effect across all sites and to assess whether observed differences in diversity were statistically significant at both local and regional scales.

All analyses were conducted in R version 4.2.2 (R Core Team, 2022). Diversity metrics were calculated using the packages *iNEXT.3D* (Chao et al., 2021, 2023) and *iNEXT.beta3D* (Chao et al., 2023). Trait distance matrices were generated using the Gower metric implemented in the *cluster* package (Maechler et al., 2022).

We then generated forest plots to visualize effect sizes and confidence intervals for each diversity metric using the *forestplot* package (Gordon & Lumley, 2024). This approach allowed for a clear comparison of treatment differences at the site level, as well as a summary meta-analysis across all sites.

To test whether the ESBC effects on the different diversity measures were correlated with (a) baseline environmental conditions or (b) ESBC-induced changes in environmental conditions, we used the mean values of environmental variables from the control district per forest site (a) and the site-level differences between control and ESBC districts (b). We did not include ESBC means separately, as these are already represented in the difference values. As the ESBC effect size on diversity, we extracted the site-level differences from the iNEXT.beta3D results. This resulted in eight values per diversity measure, environmental mean, and environmental effect size (one per forest site). We fitted linear models only for those diversity measures for which the meta-analyses showed significant effects. One model was fitted per diversity effect size and environmental variable (baseline and effect size). The environmental variables included: air temperature, soil pH, soil C:N ratio, soil microbial biomass, understory biomass, understory species richness, soil nitrogen, soil carbon, deadwood volume, litter depth (cm), and sand content. For sand content, we only tested if the mean sand content of the control patches had an effect on the diversity sizes, and not the sand effect size, as the ESBC treatment should not change sand content.

To further assess the robustness of our results, we fitted robust regression models using the rlm() function from the *MASS* package (Venables & Ripley, 2002), applied only to those models in which we had previously found significant correlations between nematode diversity effect sizes and environmental variables. This approach allowed us to identify potential outliers among the eight forest sites and to evaluate whether the regression relationships differed substantially from those obtained with the ordinary least squares linear models (lm()). Given the small sample size (n = 8), where single influential observations can strongly affect parameter estimates, this robustness check was particularly important. We visually compared the regression lines from both rlm and lm by plotting them together in the same graph to assess any deviations potentially caused by influential data points.

Linear models were fitted using the function lm() from the *stats* package (R Core Team, 2022). P-values were obtained using the function anova() from the same package. Post-hoc tests were conducted using the *emmeans* package (Lenth, 2021), and compact letter displays were derived using the package *multcomp* (Hothorn et al., 2008). Predicted values (marginal effects) were computed using the package *ggeffects* (Lüdecke, 2018). All figures, except for the forest plots in the supplementary material, were created using *ggplot2* (Wickham, 2016).

## Results

We found nematodes in all soil samples, and the densities ranged from 116 nematodes (Control patch in U03) to 9,409 nematodes per 100 g dry soil (ESBC patch in B07). Nematode densities differed between forest sites, but not between the control and ESBC district in any of the forest sites (Figure 1).

### H1: Enhancing structural β complexity in forests increases both functional and taxonomic β-diversity by creating more distinct habitat types, which in turn leads to higher γ-diversity

The meta-analysis results investigating the effect of the enhancement of structural β complexity on nematode diversity across all investigated eight forest sites showed an overall lower effect on taxonomic diversity than on functional diversity. Out of nine tested diversity measures per diversity facet, taxonomic diversity significantly increased for three measures and functional diversity decreased for four and increased for two measures (Figure 2).

**Figure 2:**
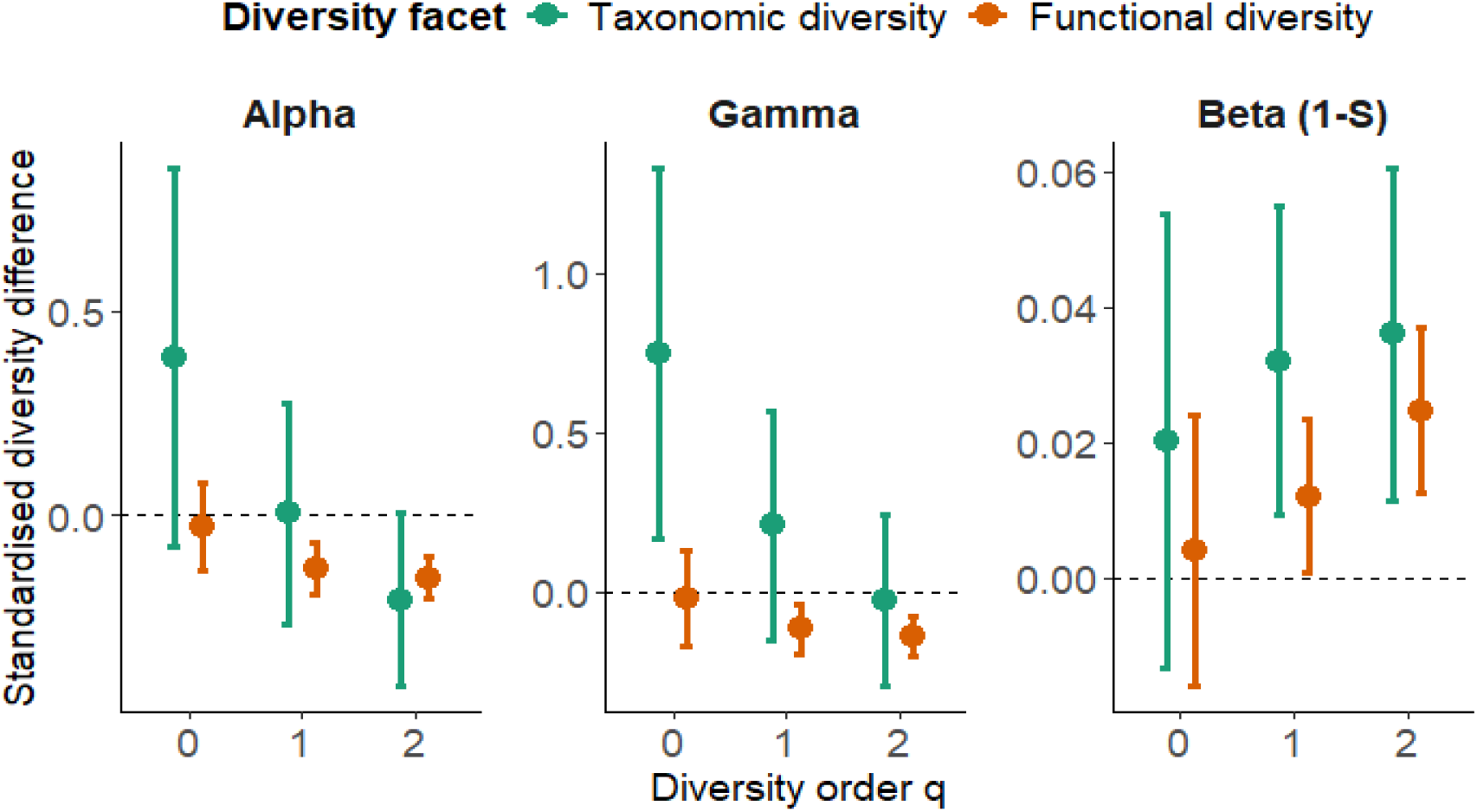
Meta-analysis results across eight forests. Points represent predicted mean values, and error bars show the 95% confidence intervals at a fixed coverage level of 0.849 (Cmax). The standardized diversity difference is calculated between each pair of control and treatment (ESBC) district for two diversity facets (taxonomic and functional diversity) and three diversity orders (q = 0, q = 1, and q = 2). To account for varying numbers of sampled assemblages (i.e., 9 or 15), a Jaccard-type turnover transformation (1 – S) of multiplicative β-diversity was applied. Positive values indicate increased diversity from control to structurally enhanced forests, while negative values indicate a decrease. Effects are considered statistically significant when the 95% confidence interval does not overlap zero.

For taxonomic diversity, ESBC led to no significant changes in α-diversity of any order q. γ-diversity of rare taxa (q = 0) increased significantly by 0.75 effective species, while the other orders q did not change. However, individual effects on γ-diversity (q = 0) in forest sites ranged from significantly negative in U03 to significantly positive in B07 and B05 (range from – 4.97 to + 4.4.2; Figure 3). For β-diversity, ESBC did not affect rare (q = 0) taxa, while it increased the diversity of common (q = 1) taxa by 0.03 and frequent (q = 2) taxa by 0.04. For both diversity orders (q = 1 and q = 2), these effects were driven by two of the eight forest sites (U03, L11) that showed a significant increase in ESBC districts, while the remaining sites exhibited no significant changes (q = 1 range from –0.06 to +0.12; q = 2 range from –0.03 to + 0.26; Figure 3).

**Figure 3:**
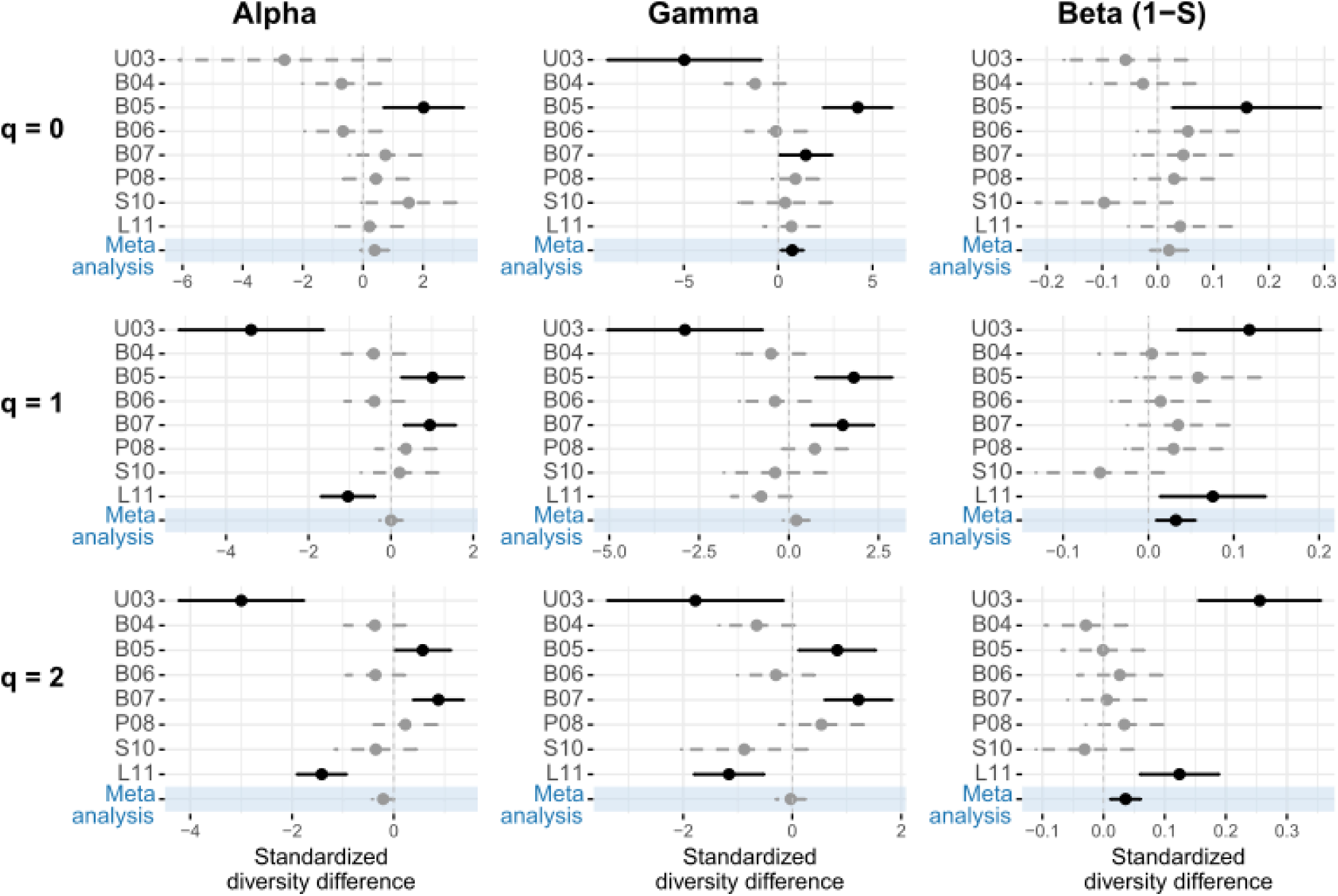
Taxonomic diversity results for individual sites and meta-analysis (across sites) at coverage level 0.849 (Cmax) with 95% confidence intervals. Standardized diversity differences between control and treatment (enhancement of structural β complexity) districts are shown for ɑ, γ, and β diversity across diversity orders q = 0, 1, and 2. A Jaccard-type turnover transformation (1 – S) was used to account for differing sample sizes. Positive values indicate higher diversity in enhanced forests; negative values indicate decreases. Effects with confidence intervals excluding zero are significant (black), while non-significant effects are grey dashed. Results of meta analyses are highlighted in blue. Individual full forest plots are in the Supporting information (Figure S5-S7).

For functional diversity, ESBC did not significantly change the α-diversity of rare (q = 0) functional taxa equivalents, but decreased the α-diversity of common (q = 1) ones by 0.13 (range from –0.86 to +0.13) and frequent (q = 2) ones by 0.16 (range from –0.88 to +0.11). The same pattern as for functional α-diversity was also found for functional γ-diversity (no change for q = 0, decreases for q = 1 and q = 2), with decreases of 0.12 (q = 1, range from –0.47 to +0.08) and 0.14 (q = 2, range from –0.43 to 0.07) functional species equivalents. Across both spatial scales (α and γ) and diversity orders (q = 1 and q =2), four forest sites (U03, B04, B06, L11) showed a significant decrease from control to ESBC, while the other four remained unchanged (Figure 4). β-diversity of rare (q = 0) functional taxa equivalents did not change significantly in response to ESBC, while β-diversity of common (q = 1) functional taxa equivalents increased by 0.01 (range from –0.03 to +0.13) and frequent ones by 0.03 (range from –0.06 to +0.18). For q = 1, significant increases were found in three forest sites (U03, P08, L11), and no change in the others. For q = 2, the same three sites (U03, P08, L11) showed significant increases, while one site (B07) showed a significant decrease (Figure 4).

**Figure 4:**
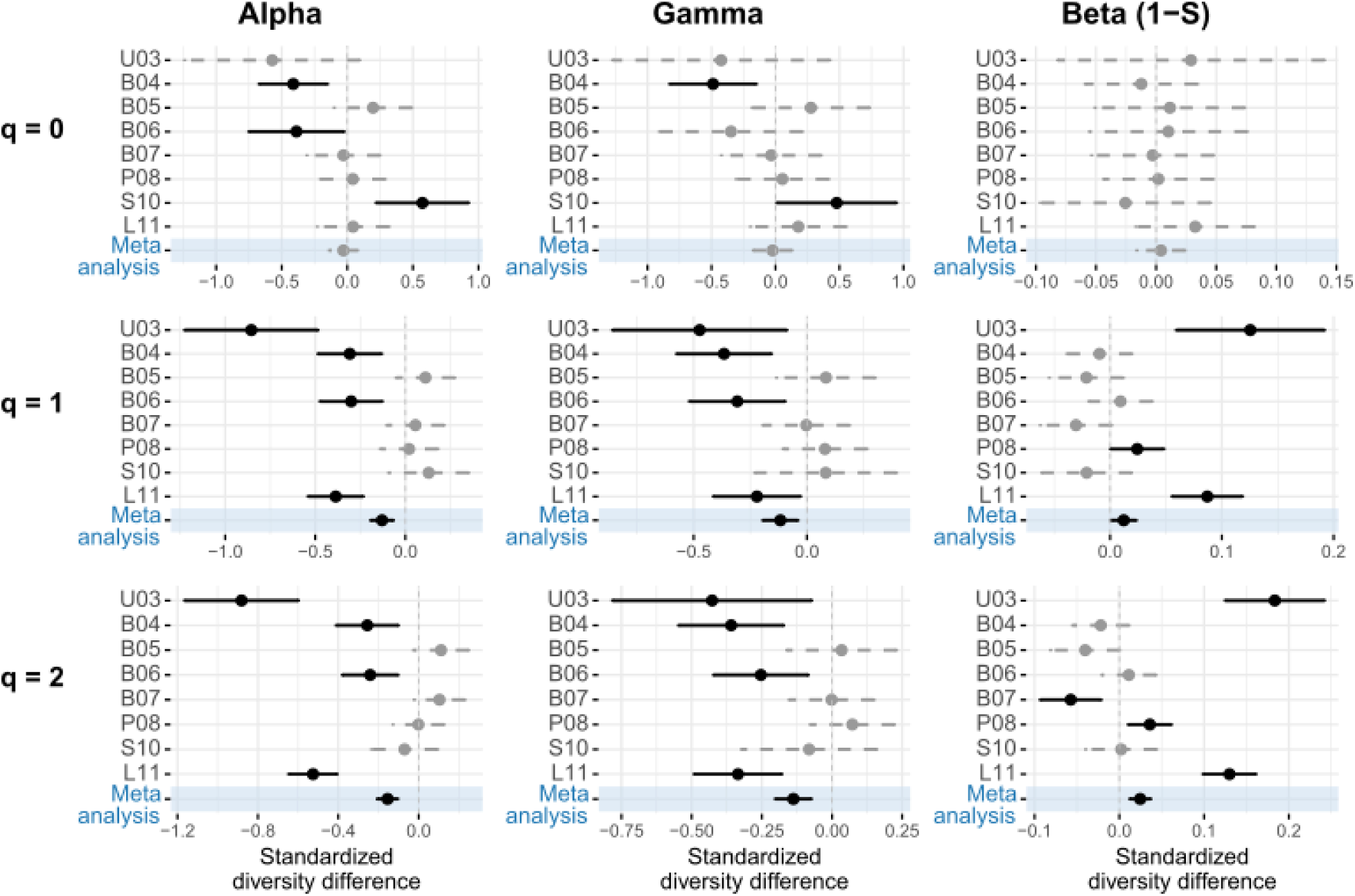
Functional diversity results for individual sites and meta-analysis (across sites) at coverage level 0.849 (Cmax) with 95% confidence intervals. Standardized diversity differences between control and treatment (enhancement of structural β complexity) districts are shown for ɑ, γ, and β diversity across diversity orders q = 0, 1, and 2. A Jaccard-type turnover transformation (1 – S) was used to account for differing sample sizes. Positive values indicate higher diversity in enhanced forests; negative values indicate decreases. Effects with confidence intervals excluding zero are significant (black), while non-significant effects are grey dashed. Results of meta analyses are highlighted in blue. Individual full forest plots are in the Supporting Information (Figure S8-S10).

The results were robust across different levels of sample coverage. Analyses with higher sample coverage (0.974) showed a similar pattern for TD: β-diversity results stay the same, while γ-diversity q = 0 was not significantly increased, and α-diversity q = 2 was significantly decreased (Figure S2 and S3 in Supporting Information). For FD, the same pattern was observed with both sample coverages (Figure S2 and S4 in Supporting Information).

### H2: Changes of nematode diversities correlate with changes of environmental variables

Out of 189 linear models testing the relationship between nematode diversity effect sizes due to ESBC and environmental variables (mean and effect sizes), 30 were significant (Table 1, Figure S11-S16 in Supporting Information).

**Table 1:**
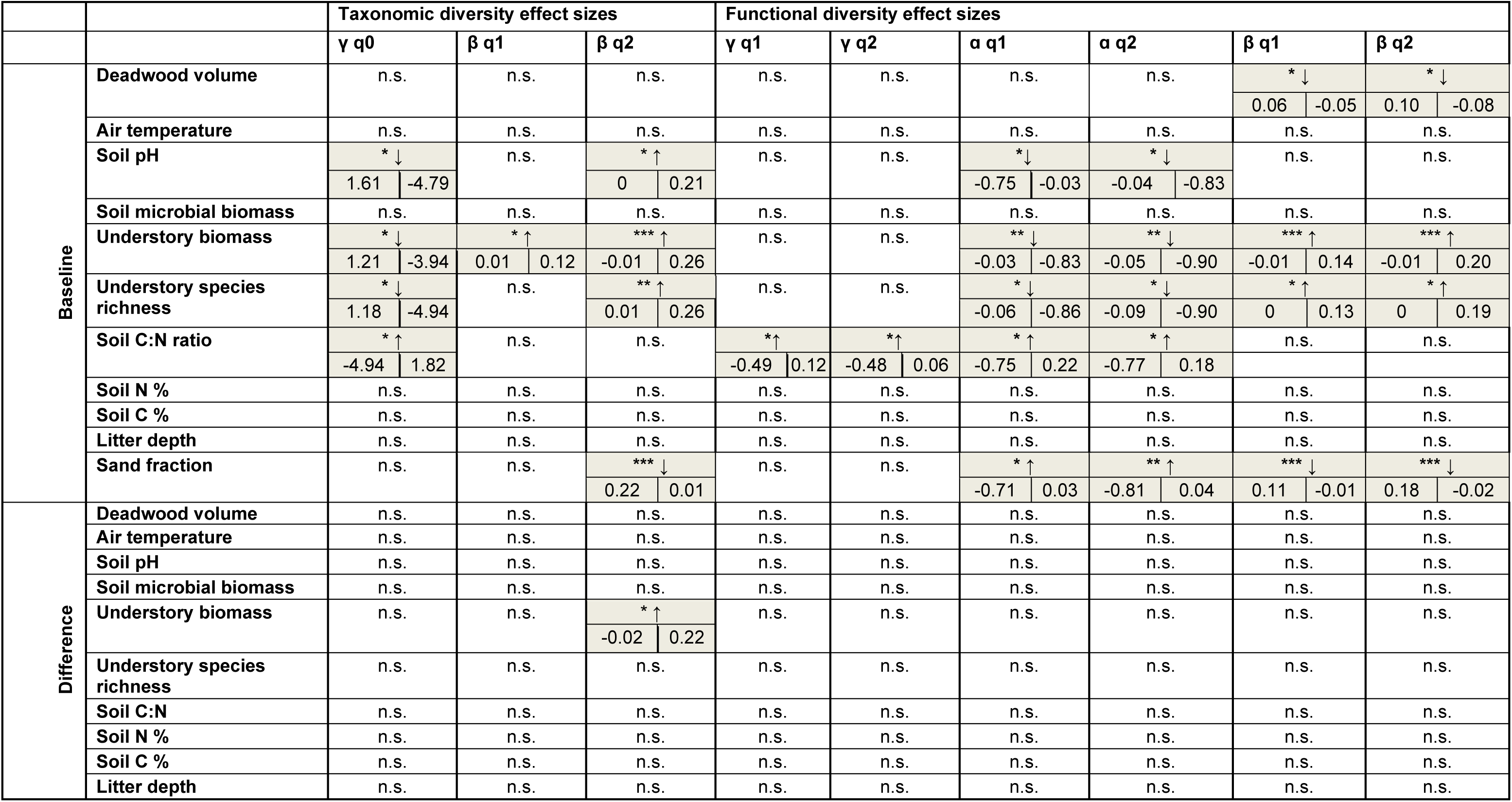
Linear regressions testing correlations between nematode diversity effect sizes (due to enhancement of structural β complexity = ESBC) and environmental variables. The table shows baseline environmental values per forest site (upper half) and corresponding ESBC effect sizes (lower half). Significant correlations are highlighted in grey. Asterisks indicate significance (p < 0.05, ** p < 0.01, *** p < 0.001) with arrows denoting regression direction. For significant results, predicted effect size ranges are shown, oriented by relationship direction (negative: larger value first, positive: smaller value first). n.s. = non-significant.

The mean values of environmental variables per forest site showed varying relationships with diversity effect sizes associated with ESBC (Table 1, Figures S5 in Supporting Information). The effect size of taxonomic diversity (TD) at the γ level for q = 0 became increasingly negative with increasing pH, higher understory biomass, and greater understory species richness. The effect size was most negative at low C/N ratios and approached zero or became positive at higher C/N ratios (Figure S11-S16 in Supporting Information).

The effect size of TD β q = 1 increased with higher understory biomass. For TD β q = 2, effect sizes increased with higher pH, greater understory biomass, higher understory species richness, and stronger ESBC effects of the understory vegetation. In contrast, the ESBC effect on TD β q = 2 declined and approached zero with increasing sand content.

The effect size of functional diversity (FD) at the γ level (q = 1 and q = 2) became less negative with increasing C/N ratio, but did not correlate with other environmental variables. The effect size of FD α q = 1 became more negative with increasing soil pH, understory biomass, and understory species richness. A decreasing ESBC effect on FD α q = 1 became less pronounced with higher C/N ratio and greater sand content. Effect sizes of FD α q = 2 became less negative with increasing C/N ratio and sand content, while the ESBC effect on FD α q = 2 became more negative with increasing pH, understory biomass, and understory species richness.

The ESBC effect on FD β q = 1 was weaker at sites with higher deadwood volume and higher sand content, but became stronger with increasing understory biomass and understory species richness. For FD β q = 2, the effect weakened with higher deadwood volume and sand content, while increasing understory biomass and species richness strengthened the effect.

Mean values of air temperature, N%, C%, and litter depth did not show any significant correlations with any diversity effect sizes. Similarly, the effect sizes of environmental variables themselves did not correlate significantly with any diversity effect size—except for understory biomass, whose effect size correlated positively with TD β q = 2. Figures of significant correlations are in the Supporting information (Figure S11-S16 in Supporting Information)

Comparison of the regression lines from robust regression models and standard linear models showed that results were largely consistent across all response–predictor combinations. Visual inspection of the 30 plots revealed that potential outliers had only minor effects on model outcomes, as robust and standard regression lines overlapped closely in most cases.

In the robust regression models, all sites were down-weighted, though with varying frequency across the 30 models (B04: 3 times, B05: 5 times, B06: 2 times, B07: 1 time, L11: 8 times, P08: 3 times, S10: 6 times, U03: 2 times). While L11 was down-weighted most often, suggesting it deviated most strongly from the overall pattern, the influence of individual sites varied depending on the specific response and predictor, and no systematic pattern of site influence was evident (Table 2S).

## Discussion

Our results demonstrate that enhanced structural β complexity (ESBC) in managed German forests does not significantly affect nematode density, but does influence various measures of nematode taxonomic and functional diversity across α-, β-, and γ-diversity scales. While some environmental variables were associated with diversity patterns, no consistent relationships emerged across all scales or diversity dimensions. Nevertheless, results indicate that initial environmental conditions impact ESBC effects on nematode diversity.

### Diversity changes of nematodes

Our first hypothesis was only partially supported, as the effects of ESBC on nematode diversity were not consistent across diversity scales, Hill numbers, and between taxonomic and functional diversity. A greater number of functional diversity measures showed significant responses compared to taxonomic diversity. This may reflect that environmental changes, such as increased habitat heterogeneity, more consistently influence the distribution of functional traits than taxa identities (Hu et al., 2014). In structurally heterogeneous forests, shifts in microhabitats and resource availability can favor or exclude species with certain ecological strategies, even if species richness remains stable (Heidrich et al., 2023; Mori et al., 2015). This underscores the value of nematode guilds—defined by trophic group and cp value and in this study complemented by body mass—as an ecologically meaningful framework for capturing such trait-based shifts (Bongers & Bongers, 1998; Cesarz et al., 2015). As functional diversity is more closely linked to ecosystem functioning than taxonomic diversity (Gagic et al., 2015), these different responses highlight the importance of considering both diversity dimensions. Our findings suggest that relying solely on taxonomic diversity may underestimate the ecological consequences of enhanced structural heterogeneity in forest ecosystems. Taxonomic γ-diversity of rare species was higher in ESBC forests, indicating more rare species at the forest level and suggesting a greater variety of niches across the forest. Interestingly, no significant effect on α or β-diversity of rare species was found. Instead, taxonomic β-diversity increased for more frequent and common species, even though their α and γ-diversity remained unchanged. Since β-diversity is derived from the ratio of γ to α, this increase must result from subtle shifts that add up to a significant change of the ratio. The observed decline in both functional α– and γ-diversity for frequent and common species equivalents in ESBC forests, combined with an increase in functional β-diversity, suggests a functional shift in the nematode communities. This may reflect a transition from generalist-dominance and broad species coexistence in homogeneous forests to stronger environmental filtering and local niche specialization in structurally complex, heterogeneous forests. This can lead to each patch hosting a more distinct set of functional traits, increasing dissimilarity between local communities (β). This in turn, can enhance multifunctionality at the landscape scale (Grman et al., 2018; van der Plas et al., 2016). While enhancing β-diversity was an intended outcome of increasing structural β complexity in forests (Müller et al., 2023), this effect was achieved at the expense of α– and γ-diversity. This means that dissimilarity between local communities increased not because more species were added, but because fewer species now occur across sites, reducing diversity at both patch and landscape scales (Figure 5). Such a pattern parallels findings from fragmented landscapes, where species turnover increases but fails to offset overall biodiversity loss (Gonçalves-Souza et al., 2025). Consequently, maintaining a mosaic of both structurally complex and more homogeneous patches may be necessary to balance local filtering effects and support broader landscape-level diversity (Eisenhauer et al., 2023).

**Figure 5:**
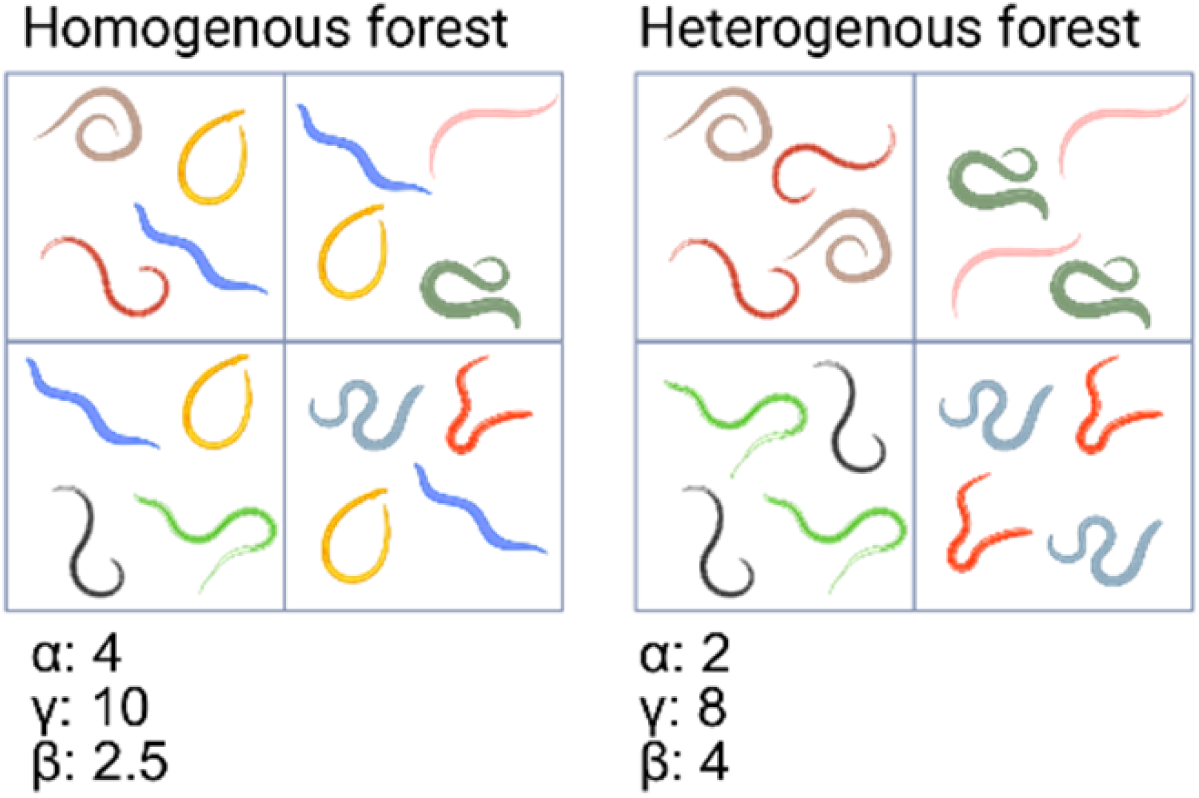
Conceptual figure of changes in nematode diversity from a homogenous forest to a heterogenous forest with enhanced structural β complexity (ESBC). In homogeneous forests, both generalist and specialist species contribute to α– and γ-diversity, with some species shared across patches, resulting in low β-diversity. In contrast, heterogeneous forests (ESBC) support more distinct communities at the patch level. This leads to decreased α– and γ-diversity (fewer shared species), but increased β-diversity. Figure designed using bioRender.

Although patterns were statistically significant, the changes in nematode diversity were rather small. Therefore, the ecological relevance may be limited and should be interpreted with caution. Nevertheless, we did find significant changes in nematode communities, which shows that community composition is indeed changing in response to aboveground structural interventions. Importantly, the observed variation across different dominance structures (Hill numbers) and scales (α, β, γ), underscores the need for a multidimensional approach. Relying on a single index (e.g., species richness alone) or a single scale could have led to different or even misleading conclusions (Chase et al., 2020; Magurran, 2021; Roswell et al., 2021).

Our meta-analytical approach revealed variable effects across sites for the same diversity measures, particularly for taxonomic diversity, but also for functional diversity (TD α and γ, q = 1, q = 2; FD α q = 0). This variability across forest sites not only explains the generally small effect sizes of the meta analyses, but also resulted in non-significant overall meta-analytical outcomes, even though three to four out of eight sites showed significant effects (Figure 3 and 4). This variability of results is in line with previous studies on the effects of forest management on nematode communities, which likewise reported varying responses (Biryol et al., 2024; Matlack, 2001; Renčo et al., 2015; Richter et al., 2023). Reported effects vary depending on forest type, soil layer, region, and the specific nematode metric examined. One study even concluded that aboveground properties, such as canopy openness, vegetation structure, litter depth, and burning, had no measurable effect on nematode communities, and that only strong soil disturbance (e.g., bedding and disking) induced significant changes (Matlack, 2001). This variability in effects shows that local site conditions strongly influence how soil nematode communities respond to aboveground structural changes, which has also been shown for soil microbial biomass and activity in the same study sites (Schwarz et al., 2025). This makes it difficult to generalize soil biodiversity responses to forest interventions. Similar to our findings for taxonomic and functional diversity across diversity scales and Hill numbers, it highlights the need for a multidimensional, multi-site approach to avoid oversimplifying results. To examine site dependency, we tested in our second hypothesis whether biodiversity responses were related to environmental variables. Despite the overall variability, the multi-site approach strengthens our conclusions for TD β (q = 1, q = 2), FD α and γ (q = 1, q = 2) as well as FD β (q = 2), because all significant site-level results for these measures pointed in the same direction. However, even within these consistent trends, effect sizes varied markedly between sites— for example, site U03 showed a 12% increase for FD β (q = 1) due to ESBC, whereas other sites showed no significant effect.

Future studies can benefit from even further expanding approaches. Examining which taxa increase or decrease, as well as applying classical nematode indices, would provide a clearer picture of community reorganization (Bongers, 1990; Du Preez et al., 2022; Ferris et al., 2001). However, such approaches were limited in our study due to the reliance on sample coverage-based extrapolation, which was necessary given the low numbers of nematodes identified in some samples. Additionally, partitioning β-diversity into turnover and nestedness components could help disentangle the underlying processes driving diversity changes (Baselga, 2010; Legendre, 2014; Montoya-Sánchez et al., 2023). To further explore biodiversity–ecosystem functioning relationships, a broader range of nematode functional traits should be integrated (Zhang et al., 2024), and diversity metrics should be directly linked to ecosystem function data or complemented with energy flux calculations (Barnes et al., 2018).

### Correlations between environmental variables and nematode diversity changes

Our findings only partially support H2, which states that environmental variables and their induced change in ESBC explain the effect sizes of nematode diversity metrics. We found correlations between nematode diversity effect sizes and environmental variables. While these patterns are complex and need to be further investigated, results indicate that initial understory vegetation biomass and sand content modulate ESBC effect on nematode diversity measures.

Baseline environmental conditions, represented by mean values from control plots, were more frequently correlated with nematode diversity effect sizes than were the effect sizes of environmental variables themselves (29 vs. 1 significant correlations). This suggests that the initial state of the environment may play a more influential role in shaping nematode communities than changes in environmental parameters induced by ESBC. This is in line with findings from a former study on the same patches that reported only minor effects of enhanced structural complexity on abiotic soil variables (Schwarz et al., 2025). Furthermore, the absence of correlations between deadwood effect sizes (*i.e.*, the experimental increase of deadwood volume) and nematode diversity effect sizes supports previous studies that found no significant influence of deadwood on nematode communities (Matlack, 2001; Renčo et al., 2015; Richter et al., 2023). Previously reported positive correlations between nematode species richness and soil nitrogen and carbon content (Renčo et al., 2015) were not observed in our dataset, even though these abiotic soil variables were the only ones that were significantly changed due to the enhancement of structural complexity (Schwarz et al., 2025). Moreover, contrary to expectations based on previous evidence that canopy gaps and the resulting understory vegetation can be important structural drivers of nematode communities (Biryol et al., 2024; Renčo et al., 2015; Thakur et al., 2014), effect sizes of understory biomass and understory species richness were also not correlated with nematode diversity effect sizes, except for one significant relationship. In our results, we found that higher baseline understory biomass and species richness were often associated with stronger treatment effects on nematode diversity, irrespective of the direction of the effect. These correlations, however, were largely driven by a single site (U03), which exhibited exceptionally high understory biomass and diversity at the outset. Comparable site-driven patterns were also apparent for soil pH, soil C/N ratio, and sand content. For vegetation biomass and sand content, site L11 also showed comparatively high baseline values and diversity effect sizes and contributed to this pattern. To assess whether patterns were disproportionately influenced by a single site, we conducted robust regression analyses. These showed that, while U03 was a site that was in many cases distinct from the others regarding environmental variables, its influence on slope estimates was generally small and did not warrant down-weighting. U03 exhibited in many cases strong ESBC effects on the nematode diversity measures (Figure 3 and 4) and combined with its distinct environmental variables, this contributed to statistically significant regression results. Even though L11 was down-weighted most frequently in the robust regression models (8 out of 30), this was not the case for models involving vegetation biomass or sand content. This supports the robustness of these regression results.

Higher baseline sand content may have reduced the magnitude of ESBC effects on nematode diversity because sandy soils have a lower water-holding capacity (Rout & Arulmozhiselvan, 2019). In such soils, any potential increase in soil moisture through canopy gaps (less trees using water) and added deadwood may be less effective, as water drains more quickly. Since nematodes depend on water films for movement (Bongers & Ferris, 1999; Neher, 2010), lower soil moisture (through higher sand content) might limit their response to environmental changes. In contrast, sites with higher initial understory biomass may have provided greater organic inputs and a more complex microhabitat from the beginning (Deng et al., 2023), thereby amplifying treatment effects on nematode diversity. These contrasting effects highlight the strong influence of regional environmental context and baseline conditions in shaping nematode community responses (Richter et al., 2023).

To reliably confirm these patterns, and particularly those related to other environmental variables, a larger number of study sites is required. Further, future research should measure soil water content over several weeks to assess the role of water availability in a more representative way, rather than relying on sand content as a proxy. Incorporating indicators, such as the bacterial-to-fungal ratio could further clarify how resource availability and microbial community structure shape nematode diversity (Hu et al., 2016). Finally, accounting for interactions among environmental variables will be key to disentangling the complex drivers of nematode community responses (Rillig et al., 2019; Thakur et al., 2018).

## Conclusion

Our study demonstrates that enhancing structural β complexity in forest ecosystems can influence soil nematode diversity. These effects are not uniform—they vary across diversity facets, scales, and dominance structures—highlighting that there is no one-size-fits-all solution to enhancing soil biodiversity. In the context of aboveground structural complexity, a mosaic of heterogeneous and homogeneous landscapes may help promote nematode diversity across scales. Evaluating multiple diversity facets and applying Hill numbers proved essential to detect changes, as more significant effects were found for functional than for taxonomic diversity, particularly in metrics sensitive to species abundance rather than richness alone. Baseline environmental factors, such as understory biomass and soil texture (sand content), can modulate nematode community responses, indicating where interventions might be most effective. This underscores the importance of considering site-specific conditions in conservation and management. To fully understand the mechanisms behind these patterns, future research should expand to more forests covering a broader range of environmental settings. Our findings provide a first step towards understanding how aboveground structural complexity can be used at forest management–relevant scales to foster more stable and multifunctional forest ecosystems through promoting soil biodiversity.

## Acknowledgements

We thank Anja Zeuner, Aron Weiss, Anne Busch, Bennet Knienieder, Felix Zeh, David Moore, Alfred Lochner, Brit Bömer, Luise Doms, Irina Kalmanova, Anne Peter, Rabea Klümpers, Clara Wild, Marina Wolz, Jens Schlüter, Josef Nußhardt, Magnus Kraatz, Anne Wendlandt, Clara Dembowski, and Christoph Stegen for their help in the field and laboratory. We also thank the entire BETA-FOR team for their support.

