## Supporting Information for "Enhanced forest heterogeneity drives stronger functional than taxonomic shifts in soil nematodes"

Table S1: Overview of all taxa (at family and genus level) recorded in this study, including their c-p value, trophic group, and average body mass (fresh weight in µg). C-p values and trophic groups were retrieved from Nemaplex on January 17, 2025, and average body mass on March 24, 2025. The c-p scale (1–5) reflects life-history strategies from colonizers (1) to persisters (5). Trophic groups: BF – bacterial feeders, FF – fungal feeders, PF – plant feeders, OM_PR – omnivores/predators.

| Taxa | c-p value | Trophic group | Average body mass [µg] |
|---|---|---|---|
| *Acrobeloides* | 2 | BF | 1.1939 |
| *Alaimus* | 4 | BF | 0.5606 |
| *Anatonchus* | 4 | OM_PR | 9.2823 |
| *Aphelenchoides* | 2 | FF | 0.1502 |
| *Aphelenchus* | 2 | FF | 0.218 |
| *Aporcelaimellus* | 5 | OM_PR | 9.0478 |
| *Aulolaimus* | 3 | BF | 0.2269 |
| *Axonchium* | 5 | PF | 2.7365 |
| *Bastiania* | 3 | BF | 0.1741 |
| *Bunonema* | 1 | BF | 0.0642 |
| Cephalobidae | 2 | BF | 0.4861 |
| *Cephalobus* | 2 | BF | 0.2662 |
| *Cervidellus* | 2 | BF | 0.1743 |
| *Chiloplacus* | 2 | BF | 0.5343 |
| *Clarkus* | 4 | OM_PR | 3.897 |
| *Coslenchus* | 2 | PF | 0.0999 |
| *Criconema* | 3 | PF | 0.6589 |
| *Criconemoides* | 3 | PF | 0.6451 |
| *Diphtherophora* | 3 | FF | 0.5039 |
| *Ditylenchus* | 2 | PF | 0.401 |
| Dolichodoridae | 3 | PF | 2.1419 |
| *Dorylaimoides* | 4 | FF | 1.1009 |
| *Ecumenicus* | 4 | OM_PR | 0.6347 |
| *Eucephalobus* | 2 | BF | 0.2358 |
| *Eudorylaimus* | 4 | OM_PR | 3.1351 |
| *Euteratocephalus* | 3 | BF | 0.263 |
| *Filenchus* | 2 | FF | 0.0988 |
| *Geomonhystera* | 1 | BF | 0.2831 |
| *Gracilacus* | 2 | PF | 0.037 |

| | | | |
|---|---|---|---|
| *Helicotylenchus* | 3 | PF | 0.2937 |
| *Heterocephalobus* | 2 | BF | 0.3559 |
| Hoplolaimidae | 3 | PF | 0.5748 |
| *Laimydorus* | 4 | OM_PR | 5.9012 |
| *Lelenchus* | 2 | PF | 0.0465 |
| Leptonchidae | 4 | FF | 0.9513 |
| *Longidorella* | 4 | PF | 0.595 |
| *Longidorus* | 5 | PF | 15.0798 |
| *Malenchus* | 2 | PF | 0.0798 |
| *Meloidogyne* | 3 | PF | 77.9998 |
| *Mesodorylaimus* | 5 | OM_PR | 1.2797 |
| *Metateratocephalus* | 3 | BF | 0.0732 |
| *Miconchus* | 4 | OM_PR | 4.7824 |
| Mononchidae | 4 | OM_PR | 6.3634 |
| *Mylonchulus* | 4 | OM_PR | 1.864 |
| *Ogma* | 3 | PF | 0.7147 |
| *Panagrolaimus* | 1 | BF | 0.6333 |
| *Paraphelenchus* | 2 | FF | 0.3356 |
| *Paratylenchus* | 2 | PF | 0.0512 |
| *Paraxonchium* | 4 | OM_PR | 5.5726 |
| *Plectus* | 2 | BF | 0.8873 |
| *Pratylenchus* | 3 | PF | 0.1438 |
| *Prismatolaimus* | 3 | BF | 0.3565 |
| *Prodorylaimus* | 4 | OM_PR | 5.6305 |
| Qudsianematidae | 4 | OM_PR | 2.8514 |
| Rhabditidae-dauerlarvae | 1 | BF | NA |
| *Rhabditis* | 1 | BF | 7.4997 |
| *Rotylenchus* | 3 | PF | 0.8595 |
| *Teratocephalus* | 3 | BF | 0.0842 |
| *Thonus* | 4 | OM_PR | 1.8915 |
| *Tripyla* | 3 | OM_PR | 4.7352 |
| Tylenchidae | 2 | FF | 0.1588 |
| *Tylencholaimellus* | 4 | FF | 0.7092 |
| *Tylencholaimus* | 4 | FF | 0.3974 |
| *Tylenchus* | 2 | PF | 0.3598 |
| *Wilsonema* | 2 | BF | 0.0608 |
| *Xiphinema* | 5 | PF | 5.4018 |

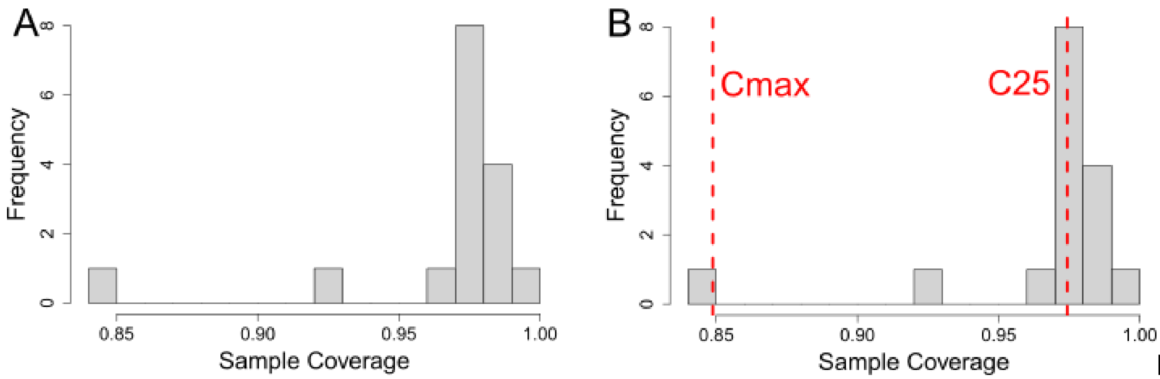


Figure S1: A) Observed (i.e., measured) sample coverages based on the original number of individuals. B) Extrapolated sample coverages assuming a doubled sample size (individual count). Red dashed lines indicate the two reference thresholds used in the analyses: Cmax, corresponding to the maximum extrapolated coverage still achievable across all sites at double sample size; and C25, representing the 25th percentile of extrapolated sample coverage values. Each unit on the y-axis is a forest site and district combination (e.g. U03 Control).

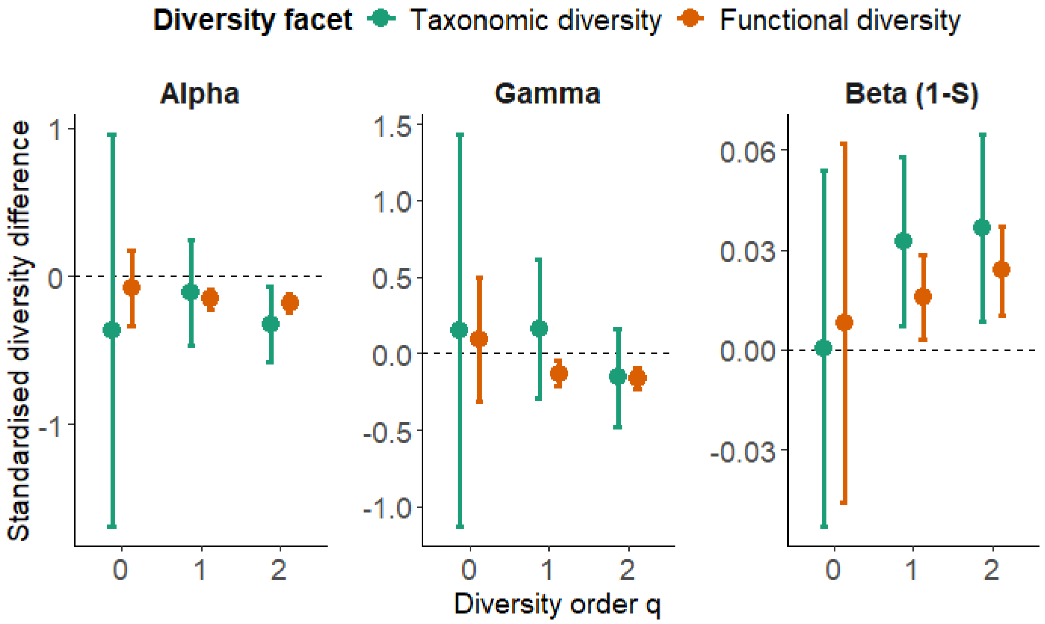


Figure S2: Meta-analysis results across eight forests. Points represent predicted mean values, and error bars show the 95% confidence intervals at a fixed coverage level of 0.974 (C25). The standardized diversity difference is calculated between each pair of control and treatment (enhancement of structural β complexity) district in directly comparable species-equivalent units for γ and α diversity and for two diversity facets (taxonomic and functional diversity) and three diversity orders (q = 0, q = 1, and q = 2).To account for varying numbers of sampled assemblages (i.e., 9 or 15), a Jaccard-type turnover transformation (1 – S) of multiplicative β diversity was applied. Positive values indicate increased diversity from control to structurally enhanced forests, while negative values indicate a decrease. Effects are considered statistically significant when the 95% confidence interval does not overlap zero.

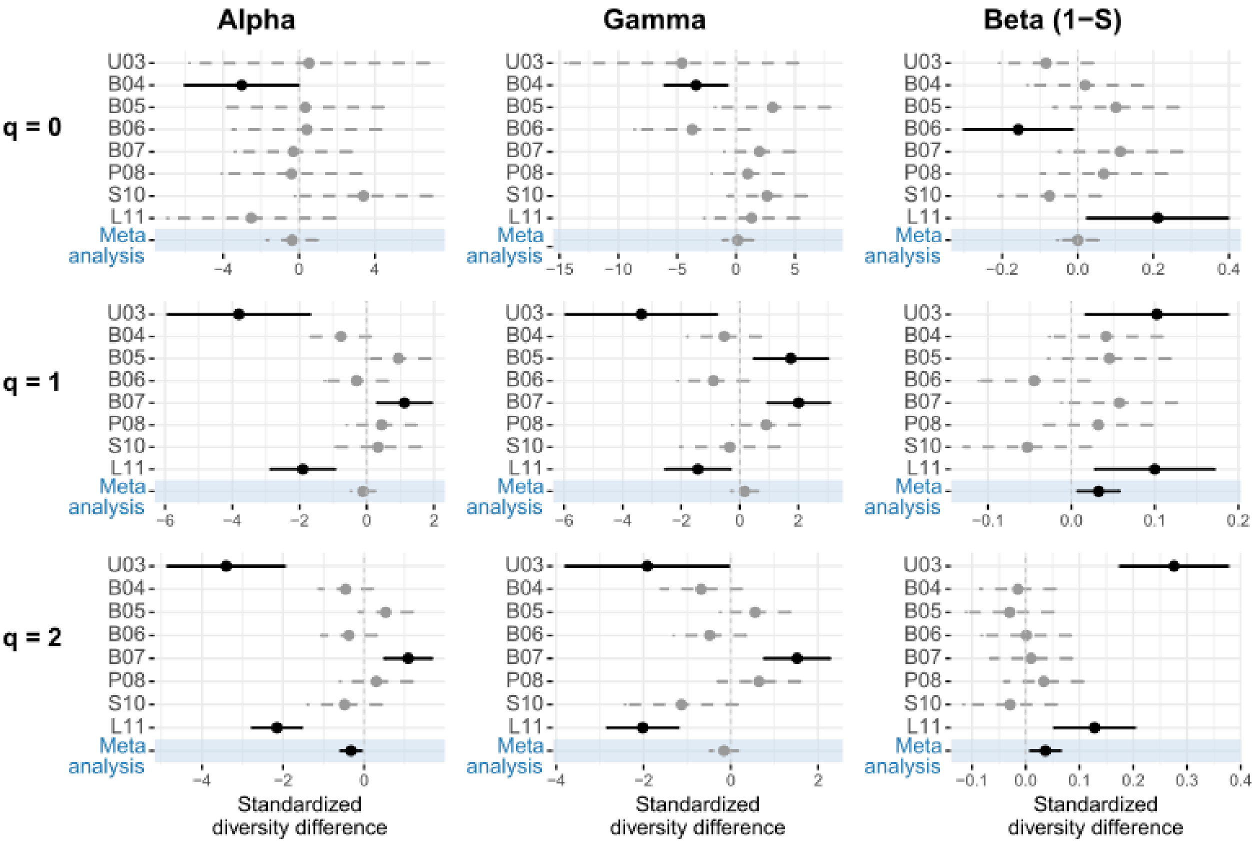


Figure S3: Taxonomic diversity results for individual sites and meta-analysis at coverage level 0.974 (C25) with 95% confidence intervals. Standardized diversity differences between control and treatment (enhancement of structural β complexity) districts are shown for α, γ, and β diversity across diversity orders q = 0, 1, and 2. A Jaccard-type turnover transformation (1 – S) was used to account for differing sample sizes. Positive values indicate higher diversity in enhanced forests; negative values indicate decreases. Effects with confidence intervals excluding zero are significant (black), while non-significant effects are grey dashed. Results of meta analyses are highlighted in blue.

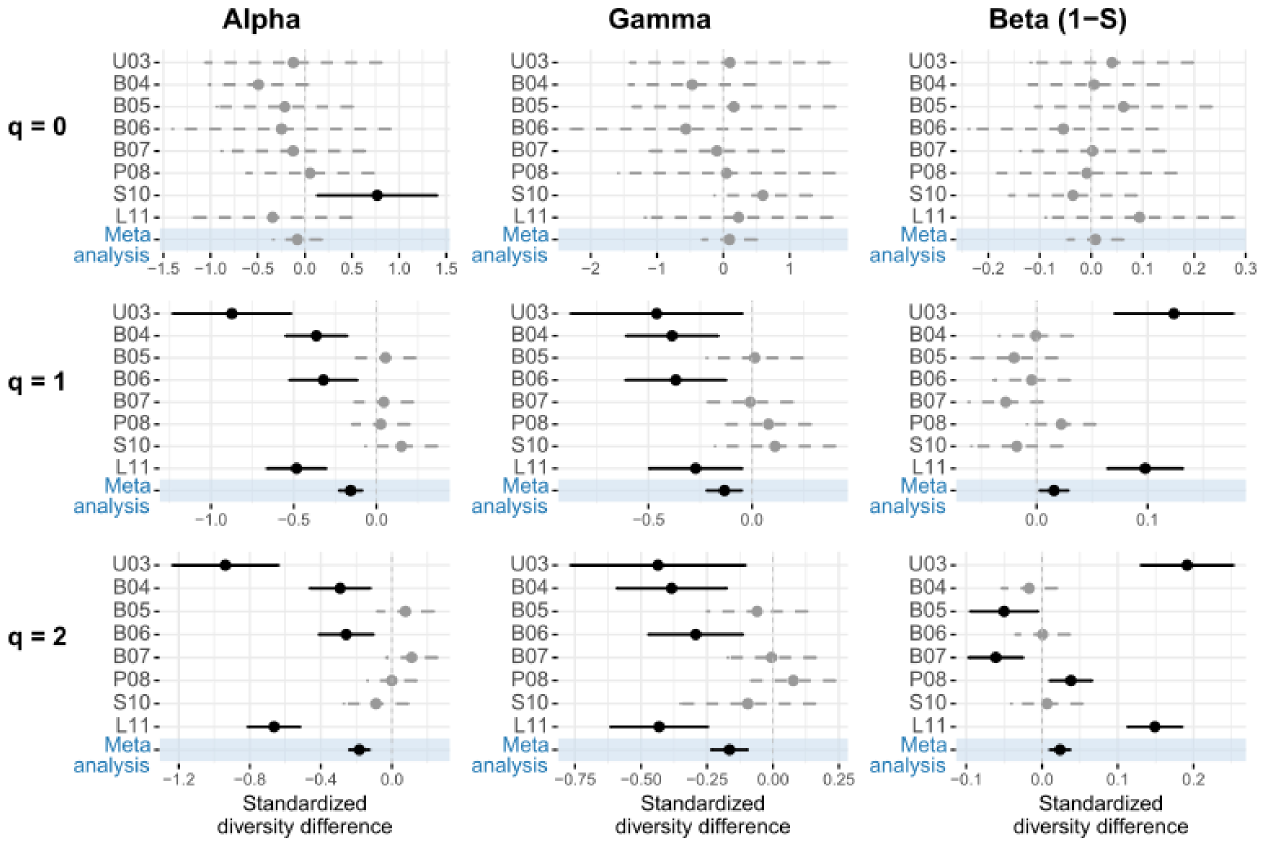


Figure S4: Functional diversity results for individual sites and meta-analysis at coverage level 0.974 (C25) with 95% confidence intervals. Standardized diversity differences between control and treatment (enhancement of structural β complexity) districts are shown for α, γ, and β diversity across diversity orders q = 0, 1, and 2. A Jaccard-type turnover transformation (1 – S) was used to account for differing sample sizes. Positive values indicate higher diversity in enhanced forests; negative values indicate decreases. Effects with confidence intervals excluding zero are significant (black), while non-significant effects are grey dashed. Results of meta analyses are highlighted in blue.

Figure S5- S7: Taxonomic diversity (TD) forest plots at sample coverage 0.849.

Individual forest plots show results per site as well as overall meta-analysis results. Plots contain values for enhanced (E; ESBC) and control (C) districts, including the difference between treatments, the lower and upper 95% confidence limits (LCL, UCL), and the fixed-effect weight per site. Each forest plot also displays the visual representation of difference with LCL and UCL for each site and for the meta-analysis. If the confidence interval does not cross zero, the effect is significant.

**A**

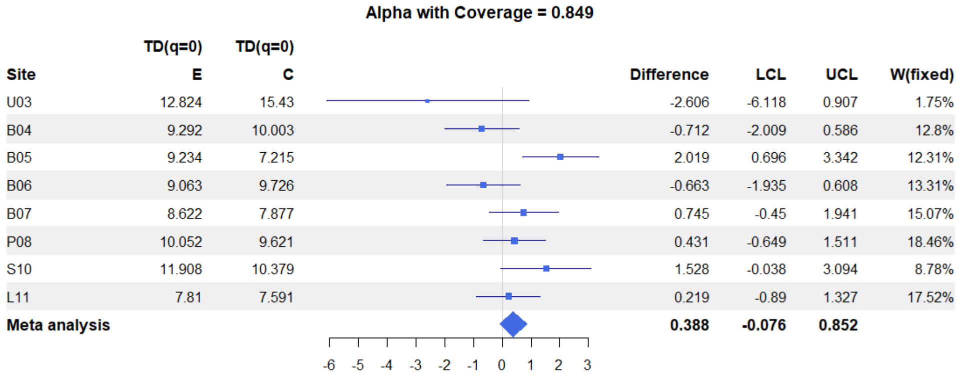


| Site | TD(q=0) E | TD(q=0) C | Difference | LCL | UCL | W(fixed) |
|---|---|---|---|---|---|---|
| U03 | 12.824 | 15.43 | -2.606 | -6.118 | 0.907 | 1.75% |
| B04 | 9.292 | 10.003 | -0.712 | -2.009 | 0.586 | 12.8% |
| B05 | 9.234 | 7.215 | 2.019 | 0.696 | 3.342 | 12.31% |
| B06 | 9.063 | 9.726 | -0.663 | -1.935 | 0.608 | 13.31% |
| B07 | 8.622 | 7.877 | 0.745 | -0.45 | 1.941 | 15.07% |
| P08 | 10.052 | 9.621 | 0.431 | -0.649 | 1.511 | 18.46% |
| S10 | 11.908 | 10.379 | 1.528 | -0.038 | 3.094 | 8.78% |
| L11 | 7.81 | 7.591 | 0.219 | -0.89 | 1.327 | 17.52% |
| **Meta analysis** | | | **0.388** | **-0.076** | **0.852** | |

**B**

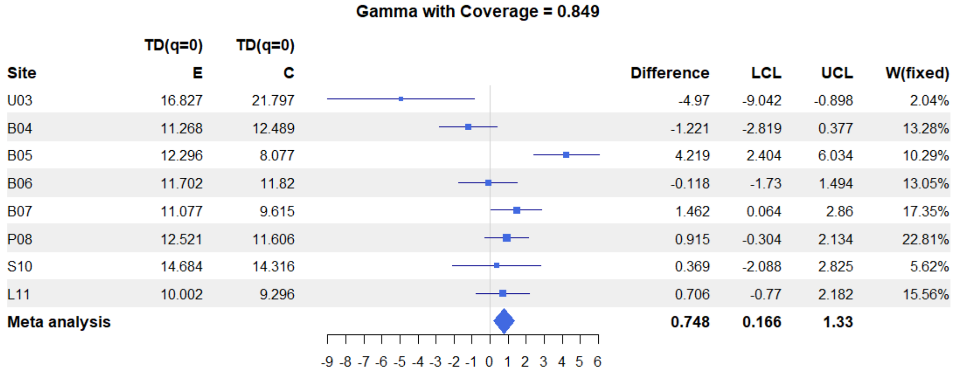


| Site | TD(q=0) E | TD(q=0) C | Difference | LCL | UCL | W(fixed) |
|---|---|---|---|---|---|---|
| U03 | 16.827 | 21.797 | -4.97 | -9.042 | -0.898 | 2.04% |
| B04 | 11.268 | 12.489 | -1.221 | -2.819 | 0.377 | 13.28% |
| B05 | 12.296 | 8.077 | 4.219 | 2.404 | 6.034 | 10.29% |
| B06 | 11.702 | 11.82 | -0.118 | -1.73 | 1.494 | 13.05% |
| B07 | 11.077 | 9.615 | 1.462 | 0.064 | 2.86 | 17.35% |
| P08 | 12.521 | 11.606 | 0.915 | -0.304 | 2.134 | 22.81% |
| S10 | 14.684 | 14.316 | 0.369 | -2.088 | 2.825 | 5.62% |
| L11 | 10.002 | 9.296 | 0.706 | -0.77 | 2.182 | 15.56% |
| **Meta analysis** | | | **0.748** | **0.166** | **1.33** | |

**C**

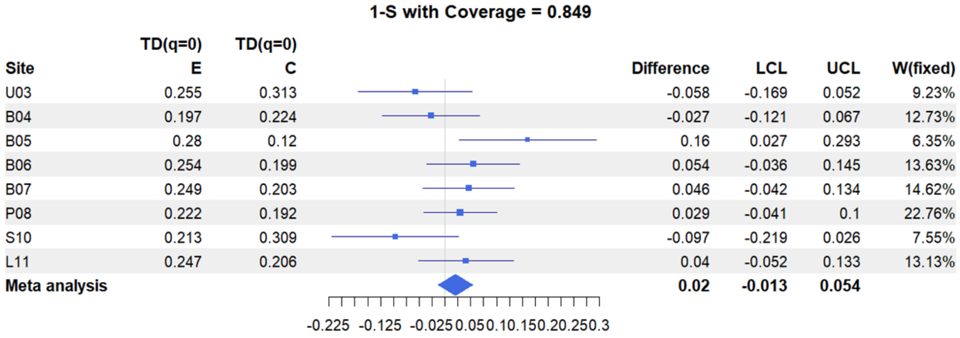


| Site | TD(q=0) E | TD(q=0) C | Difference | LCL | UCL | W(fixed) |
|---|---|---|---|---|---|---|
| U03 | 0.255 | 0.313 | -0.058 | -0.169 | 0.052 | 9.23% |
| B04 | 0.197 | 0.224 | -0.027 | -0.121 | 0.067 | 12.73% |
| B05 | 0.28 | 0.12 | 0.16 | 0.027 | 0.293 | 6.35% |
| B06 | 0.254 | 0.199 | 0.054 | -0.036 | 0.145 | 13.63% |
| B07 | 0.249 | 0.203 | 0.046 | -0.042 | 0.134 | 14.62% |
| P08 | 0.222 | 0.192 | 0.029 | -0.041 | 0.1 | 22.76% |
| S10 | 0.213 | 0.309 | -0.097 | -0.219 | 0.026 | 7.55% |
| L11 | 0.247 | 0.206 | 0.04 | -0.052 | 0.133 | 13.13% |
| **Meta analysis** | | | **0.02** | **-0.013** | **0.054** | |

Figure S5: Taxonomic diversity (TD) forest plots at coverage level 0.849. (A) α diversity, (B) γ diversity, (C) β diversity for diversity order q = 0.

**A**

**Gamma with Coverage = 0.849**

| Site | TD(q=1) E | TD(q=1) C | Difference | LCL | UCL | W(fixed) |
|---|---|---|---|---|---|---|
| U03 | 10.109 | 13.002 | -2.893 | -5.048 | -0.738 | 2.78% |
| B04 | 7.269 | 7.764 | -0.495 | -1.439 | 0.449 | 14.49% |
| B05 | 7.844 | 6.03 | 1.814 | 0.752 | 2.875 | 11.45% |
| B06 | 7.683 | 8.07 | -0.387 | -1.378 | 0.604 | 13.14% |
| B07 | 7.64 | 6.138 | 1.501 | 0.636 | 2.367 | 17.23% |
| P08 | 8.999 | 8.276 | 0.723 | -0.184 | 1.631 | 15.66% |
| S10 | 9.307 | 9.688 | -0.381 | -1.802 | 1.04 | 6.39% |
| L11 | 6.524 | 7.291 | -0.767 | -1.594 | 0.06 | 18.86% |
| **Meta analysis** | | | **0.208** | **-0.151** | **0.567** | |

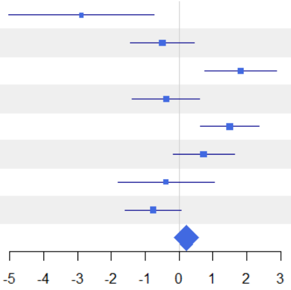


**B**

**Alpha with Coverage = 0.849**

| Site | TD(q=1) E | TD(q=1) C | Difference | LCL | UCL | W(fixed) |
|---|---|---|---|---|---|---|
| U03 | 6.844 | 10.237 | -3.393 | -5.145 | -1.641 | 2.38% |
| B04 | 5.757 | 6.175 | -0.419 | -1.182 | 0.345 | 12.54% |
| B05 | 5.72 | 4.709 | 1.011 | 0.26 | 1.761 | 12.98% |
| B06 | 5.946 | 6.348 | -0.402 | -1.12 | 0.316 | 14.18% |
| B07 | 5.782 | 4.836 | 0.947 | 0.322 | 1.572 | 18.7% |
| P08 | 7.233 | 6.868 | 0.365 | -0.373 | 1.102 | 13.45% |
| S10 | 6.791 | 6.579 | 0.211 | -0.722 | 1.144 | 8.4% |
| L11 | 4.692 | 5.733 | -1.041 | -1.69 | -0.392 | 17.37% |
| **Meta analysis** | | | **0.004** | **-0.266** | **0.274** | |

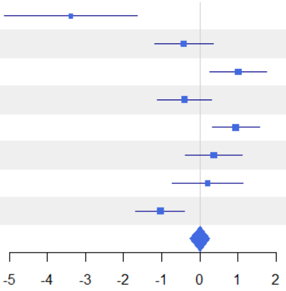


**C**

**1-S with Coverage = 0.849**

| Site | TD(q=1) E | TD(q=1) C | Difference | LCL | UCL | W(fixed) |
|---|---|---|---|---|---|---|
| U03 | 0.346 | 0.228 | 0.118 | 0.035 | 0.202 | 7.41% |
| B04 | 0.234 | 0.23 | 0.004 | -0.058 | 0.066 | 13.42% |
| B05 | 0.305 | 0.246 | 0.058 | -0.014 | 0.131 | 9.8% |
| B06 | 0.254 | 0.24 | 0.014 | -0.043 | 0.072 | 15.64% |
| B07 | 0.274 | 0.239 | 0.035 | -0.025 | 0.094 | 14.66% |
| P08 | 0.221 | 0.191 | 0.029 | -0.027 | 0.086 | 16.07% |
| S10 | 0.304 | 0.361 | -0.057 | -0.132 | 0.018 | 9.28% |
| L11 | 0.316 | 0.24 | 0.075 | 0.014 | 0.137 | 13.73% |
| **Meta analysis** | | | **0.032** | **0.009** | **0.055** | |

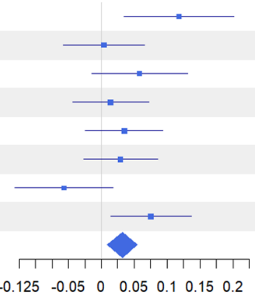


Figure S6: Taxonomic diversity (TD) forest plots at coverage level 0.849. (A) α diversity, (B) γ diversity, (C) β diversity for diversity order q = 1.

**A**

**Gamma with Coverage = 0.849**

| Site | TD(q=2) E | TD(q=2) C | Difference | LCL | UCL | W(fixed) |
|---|---|---|---|---|---|---|
| U03 | 6.487 | 8.264 | -1.777 | -3.388 | -0.166 | 2.74% |
| B04 | 4.848 | 5.5 | -0.652 | -1.341 | 0.036 | 15.02% |
| B05 | 5.708 | 4.882 | 0.826 | 0.122 | 1.531 | 14.35% |
| B06 | 5.605 | 5.904 | -0.299 | -1.005 | 0.407 | 14.28% |
| B07 | 5.605 | 4.39 | 1.215 | 0.6 | 1.831 | 18.79% |
| P08 | 6.815 | 6.281 | 0.534 | -0.237 | 1.305 | 11.99% |
| S10 | 6.349 | 7.23 | -0.88 | -2.032 | 0.271 | 5.37% |
| L11 | 4.808 | 5.968 | -1.16 | -1.799 | -0.521 | 17.44% |
| **Meta analysis** | | | **-0.028** | **-0.295** | **0.239** | |

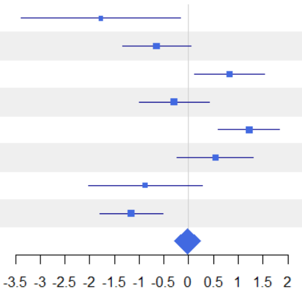


**B**

**Alpha with Coverage = 0.849**

| Site | TD(q=2) E | TD(q=2) C | Difference | LCL | UCL | W(fixed) |
|---|---|---|---|---|---|---|
| U03 | 3.747 | 6.744 | -2.997 | -4.222 | -1.771 | 2.97% |
| B04 | 3.807 | 4.179 | -0.371 | -0.968 | 0.226 | 12.54% |
| B05 | 3.893 | 3.324 | 0.568 | 0.019 | 1.118 | 14.76% |
| B06 | 4.133 | 4.493 | -0.36 | -0.938 | 0.218 | 13.35% |
| B07 | 4.148 | 3.27 | 0.878 | 0.387 | 1.37 | 18.5% |
| P08 | 5.334 | 5.105 | 0.228 | -0.391 | 0.848 | 11.62% |
| S10 | 3.991 | 4.345 | -0.355 | -1.152 | 0.443 | 7.02% |
| L11 | 3.136 | 4.551 | -1.415 | -1.897 | -0.933 | 19.25% |
| **Meta analysis** | | | **-0.208** | **-0.419** | **0.003** | |

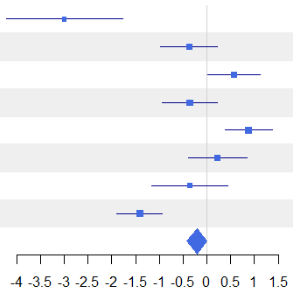


**C**

**1-S with Coverage = 0.849**

| Site | TD(q=2) E | TD(q=2) C | Difference | LCL | UCL | W(fixed) |
|---|---|---|---|---|---|---|
| U03 | 0.453 | 0.197 | 0.255 | 0.156 | 0.355 | 6.08% |
| B04 | 0.241 | 0.27 | -0.029 | -0.096 | 0.038 | 13.49% |
| B05 | 0.358 | 0.359 | -0.001 | -0.068 | 0.066 | 13.4% |
| B06 | 0.295 | 0.269 | 0.027 | -0.042 | 0.096 | 12.67% |
| B07 | 0.292 | 0.287 | 0.005 | -0.058 | 0.069 | 14.84% |
| P08 | 0.245 | 0.211 | 0.034 | -0.029 | 0.097 | 15.28% |
| S10 | 0.418 | 0.449 | -0.031 | -0.11 | 0.048 | 9.63% |
| L11 | 0.391 | 0.267 | 0.124 | 0.06 | 0.188 | 14.61% |
| **Meta analysis** | | | **0.036** | **0.011** | **0.061** | |

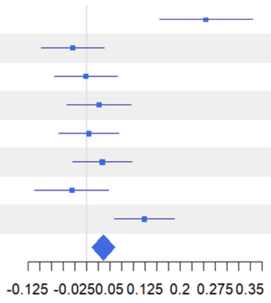


Figure S7: Taxonomic diversity (TD) forest plots at coverage level 0.849. (A) α diversity, (B) γ diversity, (C) β diversity for diversity order q = 2.

Figure S8- S10: Functional diversity (FD) forest plots at sample coverage 0.849. Individual forest plots show results per site as well as overall meta-analysis results. Plots contain values for enhanced (E; ESBC) and control (C) districts, including the difference between treatments, the lower and upper 95% confidence limits (LCL, UCL), and the fixed-effect weight per site. Each forest plot also displays the visual representation of difference with LCL and UCL for each site and for the meta-analysis. If the confidence interval does not cross zero, the effect is significant.

**A**

**Gamma with Coverage = 0.849**

| Site | FD(q=0) E | FD(q=0) C | Difference | LCL | UCL | W(fixed) |
|---|---|---|---|---|---|---|
| U03 | 4.479 | 4.903 | -0.425 | -1.273 | 0.424 | 3.14% |
| B04 | 3.148 | 3.636 | -0.488 | -0.825 | -0.151 | 19.92% |
| B05 | 3.274 | 2.997 | 0.277 | -0.181 | 0.735 | 10.79% |
| B06 | 3.049 | 3.396 | -0.348 | -0.906 | 0.21 | 7.27% |
| B07 | 2.885 | 2.918 | -0.033 | -0.423 | 0.357 | 14.9% |
| P08 | 3.045 | 2.991 | 0.054 | -0.307 | 0.415 | 17.39% |
| S10 | 4.026 | 3.547 | 0.478 | 0.011 | 0.946 | 10.37% |
| L11 | 3.121 | 2.942 | 0.179 | -0.195 | 0.552 | 16.22% |
| **Meta analysis** | | | **-0.023** | **-0.173** | **0.128** | |

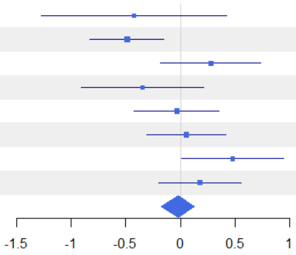


**B**

**Alpha with Coverage = 0.849**

| Site | FD(q=0) E | FD(q=0) C | Difference | LCL | UCL | W(fixed) |
|---|---|---|---|---|---|---|
| U03 | 4.413 | 4.986 | -0.573 | -1.24 | 0.095 | 2.58% |
| B04 | 3.019 | 3.43 | -0.412 | -0.671 | -0.152 | 17.04% |
| B05 | 3.07 | 2.874 | 0.196 | -0.099 | 0.491 | 13.21% |
| B06 | 2.83 | 3.218 | -0.388 | -0.751 | -0.025 | 8.73% |
| B07 | 2.722 | 2.751 | -0.03 | -0.308 | 0.248 | 14.87% |
| P08 | 2.866 | 2.824 | 0.042 | -0.204 | 0.289 | 18.93% |
| S10 | 3.8 | 3.227 | 0.573 | 0.221 | 0.925 | 9.28% |
| L11 | 2.869 | 2.825 | 0.044 | -0.229 | 0.318 | 15.36% |
| **Meta analysis** | | | **-0.029** | **-0.137** | **0.078** | |

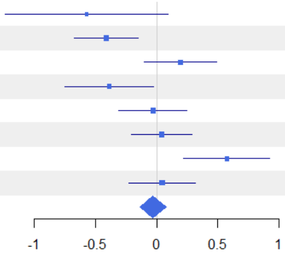


**C**

**1-S with Coverage = 0.849**

| Site | FD(q=0) E | FD(q=0) C | Difference | LCL | UCL | W(fixed) |
|---|---|---|---|---|---|---|
| U03 | 0.003 | -0.026 | 0.029 | -0.082 | 0.14 | 3.24% |
| B04 | 0.024 | 0.037 | -0.012 | -0.059 | 0.034 | 18.27% |
| B05 | 0.041 | 0.03 | 0.011 | -0.051 | 0.074 | 10.29% |
| B06 | 0.046 | 0.035 | 0.01 | -0.055 | 0.076 | 9.25% |
| B07 | 0.036 | 0.038 | -0.003 | -0.054 | 0.048 | 15.46% |
| P08 | 0.038 | 0.036 | 0.002 | -0.044 | 0.048 | 19.12% |
| S10 | 0.034 | 0.06 | -0.026 | -0.096 | 0.045 | 7.96% |
| L11 | 0.059 | 0.026 | 0.033 | -0.017 | 0.082 | 16.41% |
| **Meta analysis** | | | **0.004** | **-0.016** | **0.024** | |

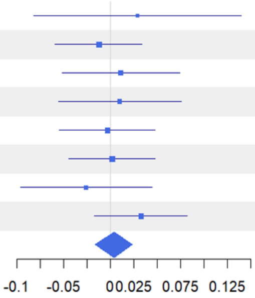


Figure S8: Functional diversity (FD) forest plots at coverage level 0.849. (A) α diversity, (B) γ diversity, (C) β diversity for diversity order q = 0.

**A**

**Gamma with Coverage = 0.849**

| Site | FD(q=1) E | FD(q=1) C | Difference | LCL | UCL | W(fixed) |
|---|---|---|---|---|---|---|
| U03 | 3.338 | 3.811 | -0.474 | -0.858 | -0.089 | 4.06% |
| B04 | 2.527 | 2.893 | -0.366 | -0.575 | -0.156 | 13.68% |
| B05 | 2.736 | 2.652 | 0.084 | -0.134 | 0.302 | 12.68% |
| B06 | 2.319 | 2.627 | -0.307 | -0.519 | -0.096 | 13.43% |
| B07 | 2.361 | 2.364 | -0.003 | -0.194 | 0.188 | 16.4% |
| P08 | 2.427 | 2.347 | 0.08 | -0.106 | 0.265 | 17.45% |
| S10 | 2.878 | 2.795 | 0.083 | -0.23 | 0.396 | 6.14% |
| L11 | 2.306 | 2.527 | -0.221 | -0.413 | -0.028 | 16.16% |
| **Meta analysis** | | | **-0.117** | **-0.194** | **-0.039** | |

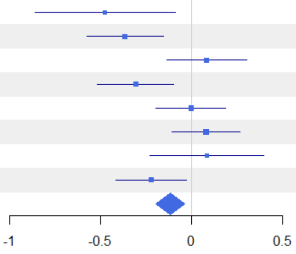


**B**

**Alpha with Coverage = 0.849**

| Site | FD(q=1) E | FD(q=1) C | Difference | LCL | UCL | W(fixed) |
|---|---|---|---|---|---|---|
| U03 | 2.721 | 3.577 | -0.855 | -1.225 | -0.485 | 2.9% |
| B04 | 2.305 | 2.614 | -0.308 | -0.485 | -0.131 | 12.64% |
| B05 | 2.324 | 2.212 | 0.112 | -0.051 | 0.274 | 15.02% |
| B06 | 2.121 | 2.422 | -0.301 | -0.474 | -0.128 | 13.25% |
| B07 | 2.176 | 2.12 | 0.056 | -0.102 | 0.215 | 15.74% |
| P08 | 2.255 | 2.234 | 0.021 | -0.139 | 0.181 | 15.5% |
| S10 | 2.408 | 2.278 | 0.13 | -0.091 | 0.351 | 8.1% |
| L11 | 1.945 | 2.331 | -0.386 | -0.54 | -0.233 | 16.85% |
| **Meta analysis** | | | **-0.129** | **-0.192** | **-0.066** | |

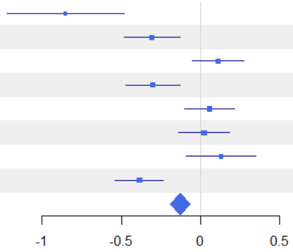


**C**

**1-S with Coverage = 0.849**

| Site | FD(q=1) E | FD(q=1) C | Difference | LCL | UCL | W(fixed) |
|---|---|---|---|---|---|---|
| U03 | 0.188 | 0.063 | 0.125 | 0.059 | 0.192 | 2.87% |
| B04 | 0.082 | 0.091 | -0.01 | -0.039 | 0.019 | 14.93% |
| B05 | 0.151 | 0.172 | -0.021 | -0.054 | 0.012 | 11.49% |
| B06 | 0.077 | 0.068 | 0.009 | -0.019 | 0.038 | 15.71% |
| B07 | 0.065 | 0.096 | -0.031 | -0.062 | 0.001 | 12.42% |
| P08 | 0.064 | 0.04 | 0.024 | 0 | 0.048 | 22.13% |
| S10 | 0.168 | 0.189 | -0.021 | -0.061 | 0.019 | 7.79% |
| L11 | 0.159 | 0.072 | 0.087 | 0.055 | 0.118 | 12.65% |
| **Meta analysis** | | | **0.012** | **0.001** | **0.023** | |

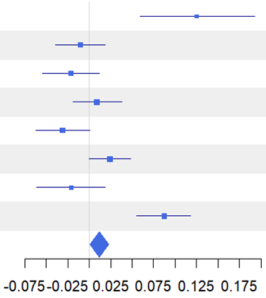


Figure S9: Functional diversity (FD) forest plots at coverage level 0.849. (A) α diversity, (B) γ diversity, (C) β diversity for diversity order q = 1.

**A**

**Gamma with Coverage = 0.849**

| Site | FD(q=2) E | FD(q=2) C | Difference | LCL | UCL | W(fixed) |
|---|---|---|---|---|---|---|
| U03 | 2.86 | 3.287 | -0.427 | -0.782 | -0.072 | 3.25% |
| B04 | 2.181 | 2.54 | -0.359 | -0.546 | -0.172 | 11.7% |
| B05 | 2.479 | 2.445 | 0.034 | -0.161 | 0.23 | 10.71% |
| B06 | 2.007 | 2.261 | -0.253 | -0.422 | -0.085 | 14.42% |
| B07 | 2.083 | 2.084 | -0.001 | -0.152 | 0.15 | 17.96% |
| P08 | 2.114 | 2.042 | 0.072 | -0.078 | 0.222 | 18.2% |
| S10 | 2.371 | 2.453 | -0.082 | -0.323 | 0.16 | 7.03% |
| L11 | 1.949 | 2.284 | -0.335 | -0.491 | -0.179 | 16.74% |
| **Meta analysis** | | | **-0.138** | **-0.202** | **-0.074** | |

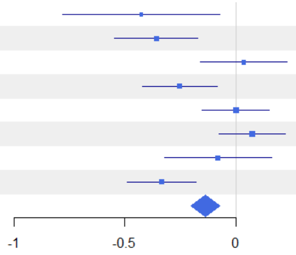


**B**

**Alpha with Coverage = 0.849**

| Site | FD(q=2) E | FD(q=2) C | Difference | LCL | UCL | W(fixed) |
|---|---|---|---|---|---|---|
| U03 | 1.981 | 2.862 | -0.881 | -1.164 | -0.598 | 3.19% |
| B04 | 1.941 | 2.197 | -0.255 | -0.41 | -0.101 | 10.71% |
| B05 | 1.955 | 1.843 | 0.112 | -0.025 | 0.249 | 13.64% |
| B06 | 1.783 | 2.025 | -0.241 | -0.379 | -0.103 | 13.34% |
| B07 | 1.89 | 1.786 | 0.104 | -0.022 | 0.229 | 16.23% |
| P08 | 1.935 | 1.936 | -0.001 | -0.128 | 0.126 | 15.84% |
| S10 | 1.772 | 1.842 | -0.07 | -0.234 | 0.094 | 9.48% |
| L11 | 1.522 | 2.048 | -0.526 | -0.646 | -0.405 | 17.57% |
| **Meta analysis** | | | **-0.155** | **-0.205** | **-0.104** | |

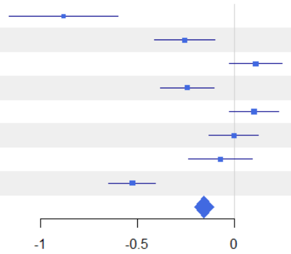


**C**

**1-S with Coverage = 0.849**

| Site | FD(q=2) E | FD(q=2) C | Difference | LCL | UCL | W(fixed) |
|---|---|---|---|---|---|---|
| U03 | 0.319 | 0.136 | 0.183 | 0.125 | 0.242 | 4.36% |
| B04 | 0.114 | 0.136 | -0.022 | -0.055 | 0.011 | 13.54% |
| B05 | 0.222 | 0.262 | -0.04 | -0.081 | 0 | 8.98% |
| B06 | 0.111 | 0.1 | 0.011 | -0.02 | 0.043 | 14.77% |
| B07 | 0.087 | 0.144 | -0.057 | -0.093 | -0.022 | 11.63% |
| P08 | 0.085 | 0.049 | 0.036 | 0.011 | 0.061 | 22.93% |
| S10 | 0.269 | 0.267 | 0.002 | -0.04 | 0.044 | 8.53% |
| L11 | 0.232 | 0.102 | 0.13 | 0.099 | 0.161 | 15.26% |
| **Meta analysis** | | | **0.025** | **0.012** | **0.037** | |

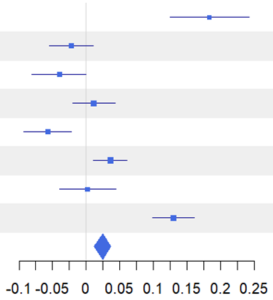


Figure S10: Functional diversity (FD) forest plots at coverage level 0.849. (A) α diversity, (B) γ diversity, (C) β diversity for diversity order q = 2.

Figure S11-S16: Regression plots of significant ($p < 0.05$) linear relationships between diversity measure effect sizes due to ESBC and environmental variables. Each plot shows the relationship between the effect size of a diversity measure and a given environmental variable. Points are colored by forest site. For each environmental variable, all significant regressions across α, γ, and β diversity and across diversity orders (q = 0, 1, 2) are presented together. Asterisks indicate the significance level of the regression (* $p \leq 0.05$, ** $p \leq 0.01$, *** $p \leq 0.001$).

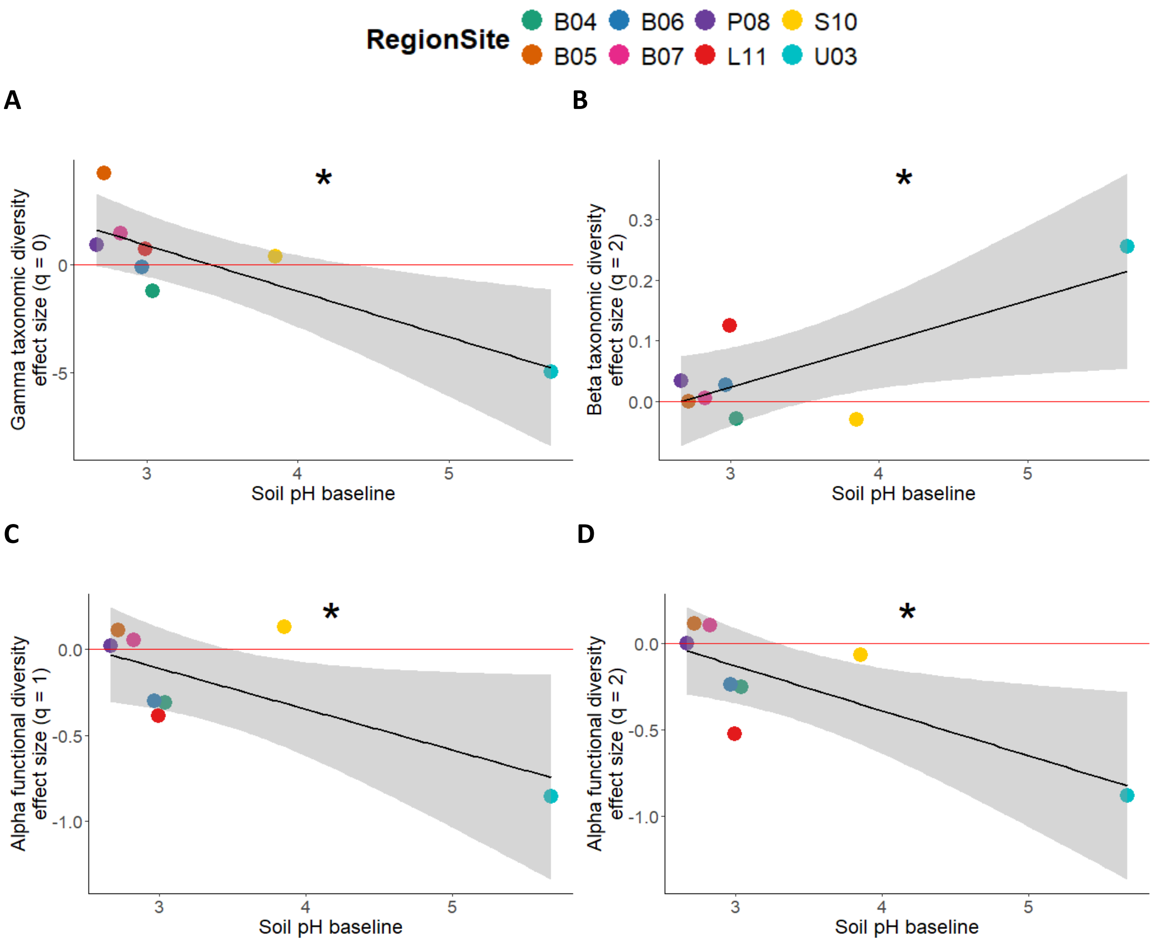


Figure S11: Regression plots of significant ($p < 0.05$) linear relationships between diversity measure effect sizes due to ESBC and basal soil pH. (A) γ taxonomic diversity, q = 0; (B) β taxonomic diversity, q = 2; (C) α functional diversity, q = 1; (D) α functional diversity, q = 2.

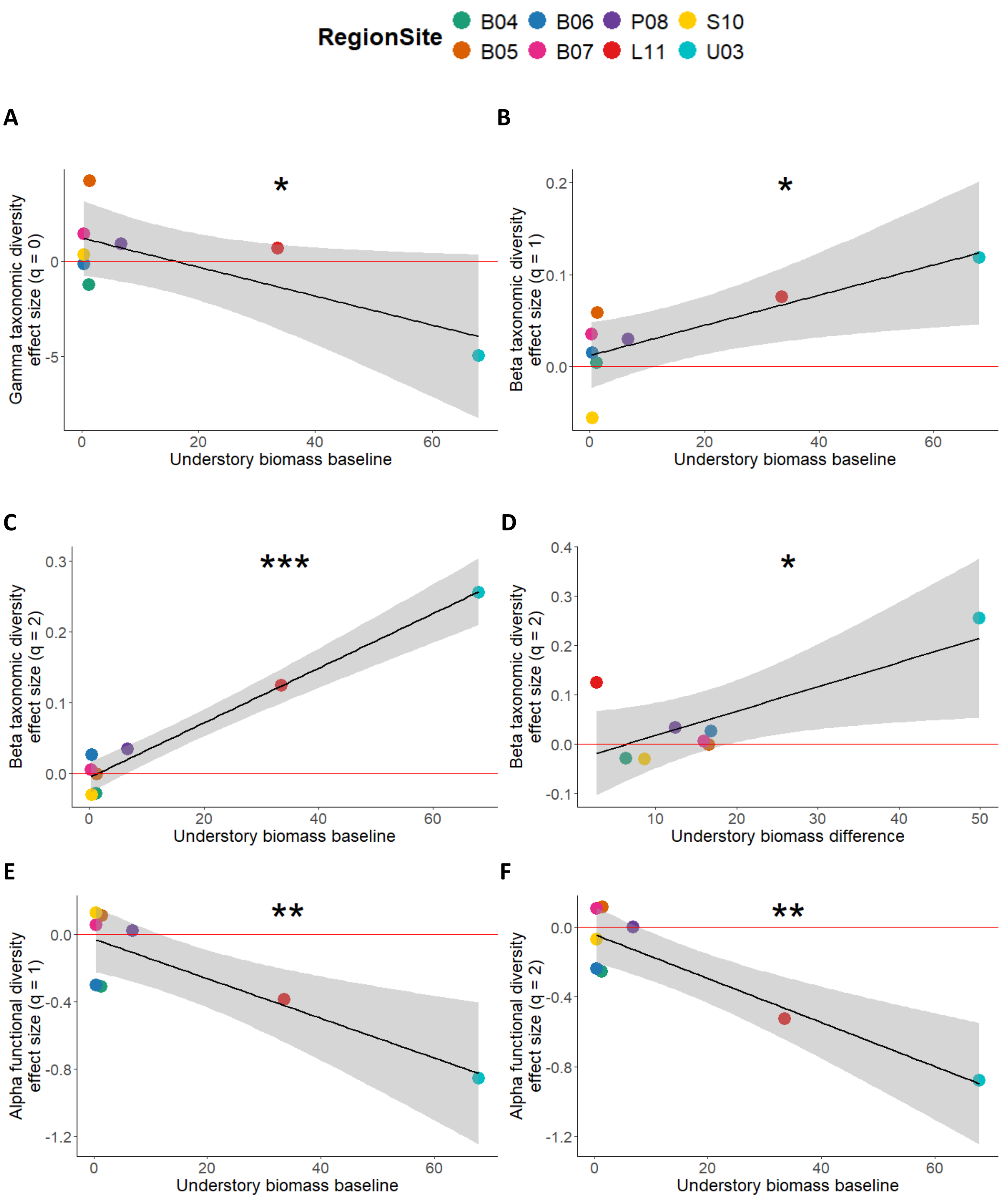
RegionSite
B04 B06 P08 S10
B05 B07 L11 U03
A
*
Gamma taxonomic diversity effect size (q = 0)
Understory biomass baseline
B
*
Beta taxonomic diversity effect size (q = 1)
Understory biomass baseline
C
***
Beta taxonomic diversity effect size (q = 2)
Understory biomass baseline
D
*
Beta taxonomic diversity effect size (q = 2)
Understory biomass difference
E
**
Alpha functional diversity effect size (q = 1)
Understory biomass baseline
F
**
Alpha functional diversity effect size (q = 2)
Understory biomass baseline

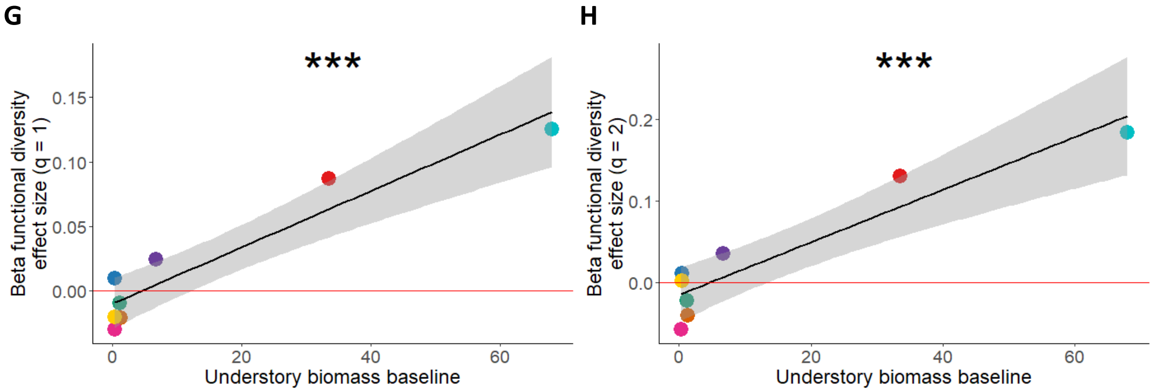


Figure S12: Regression plots of significant ($p < 0.05$) linear relationships between diversity measure effect sizes due to ESBC and understory biomass. A-C and E-F show basal understory biomass, while D shows the understory biomass effect size. (A) γ taxonomic diversity, q = 0; (B) β taxonomic diversity, q = 1; (C) β taxonomic diversity, q = 2; (D) β taxonomic diversity, q = 2; (E) α functional diversity, q = 1; (F) α functional diversity, q = 2; (G) β functional diversity, q = 1; (H) β functional diversity, q = 2.

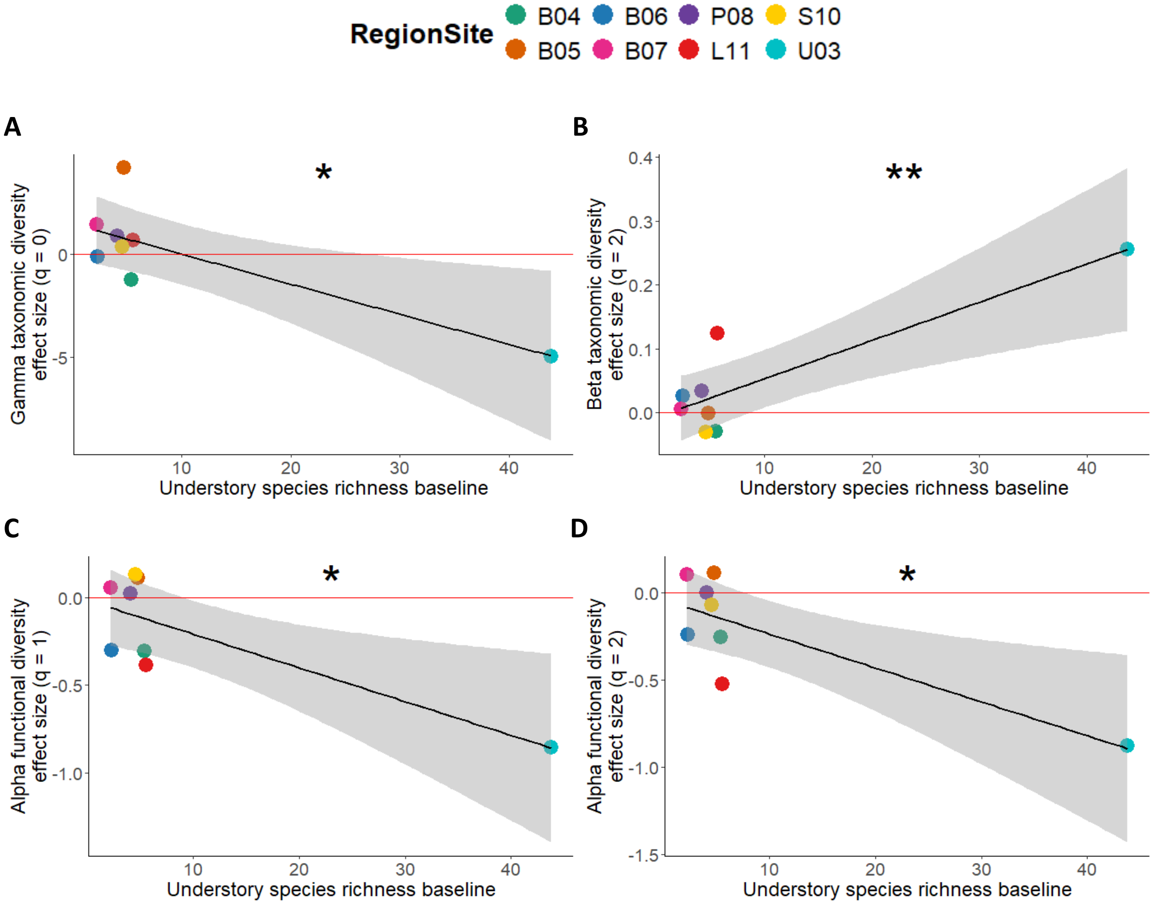

E

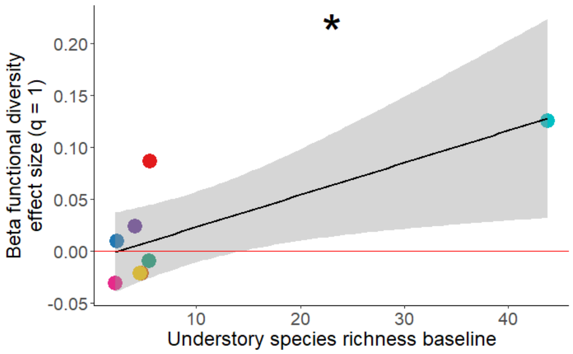


F

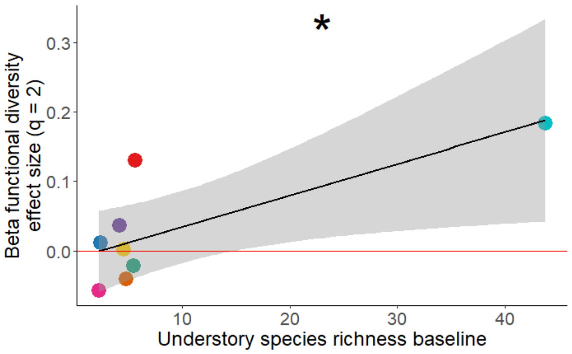


Figure S13: Regression plots of significant ($p < 0.05$) linear relationships between diversity measure effect sizes due to ESBC and understory species richness. (A) γ taxonomic diversity, q = 0; (B) β taxonomic diversity, q = 2; (C) α functional diversity, q = 1; (D) α functional diversity, q = 2; (E) β functional diversity, q = 1; (F) β functional diversity, q = 2.

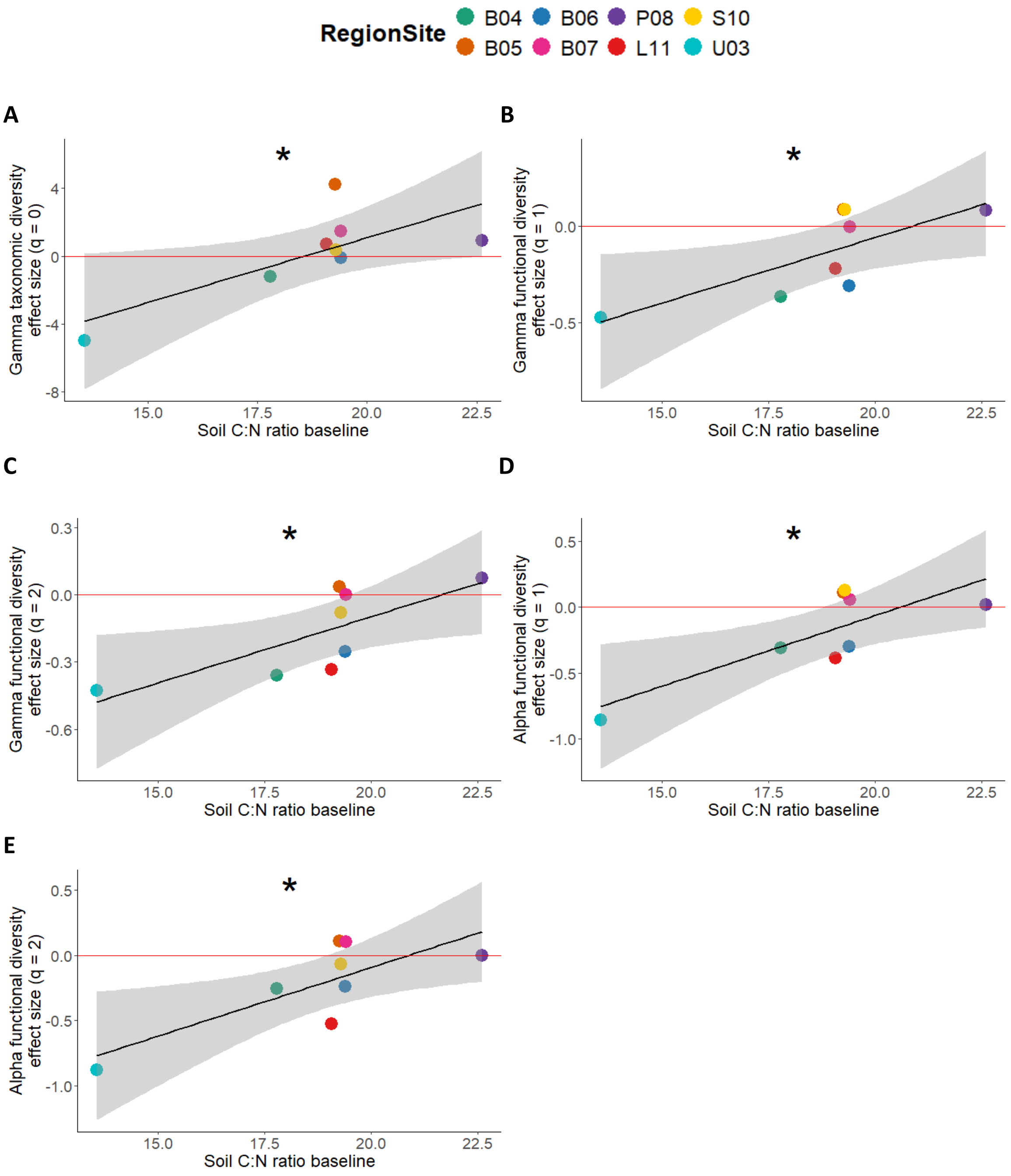


Figure S14: Regression plots of significant ($p < 0.05$) linear relationships between diversity measure effect sizes due to ESBC and soil carbon-to-nitrogen ratio (C:N). (A) γ taxonomic diversity, q = 0; (B) γ functional diversity, q = 1; (C) γ functional diversity, q = 2; (D) α functional diversity, q = 1; (E) α functional diversity, q = 2.

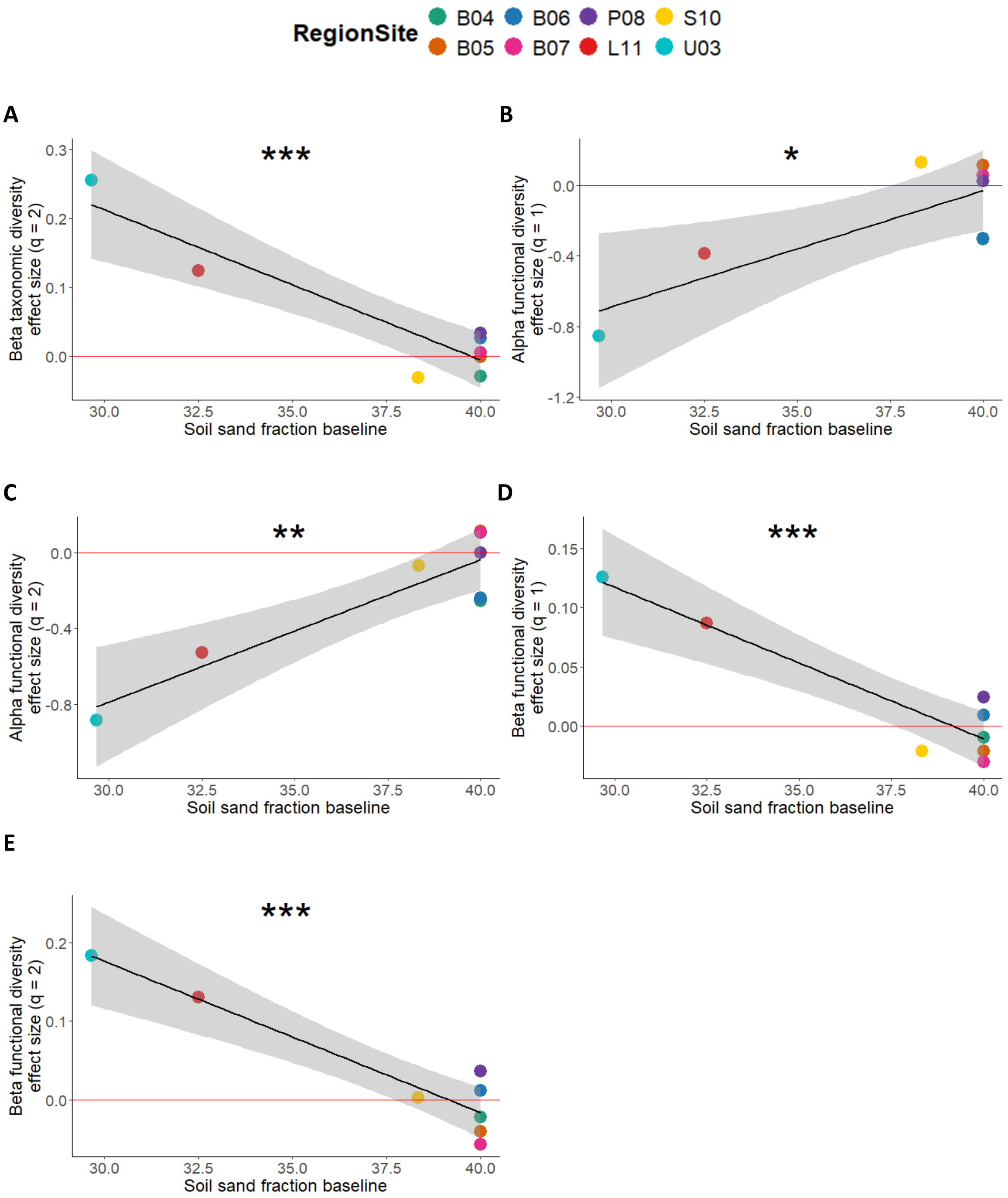


Figure S15: Regression plots of significant ($p < 0.05$) linear relationships between diversity measure effect sizes due to ESBC and soil sand content. (A) β taxonomic diversity, q = 2; (B) α functional diversity, q = 1; (C) α functional diversity, q = 2; (D) β functional diversity, q = 1; (E) β functional diversity, q = 2.

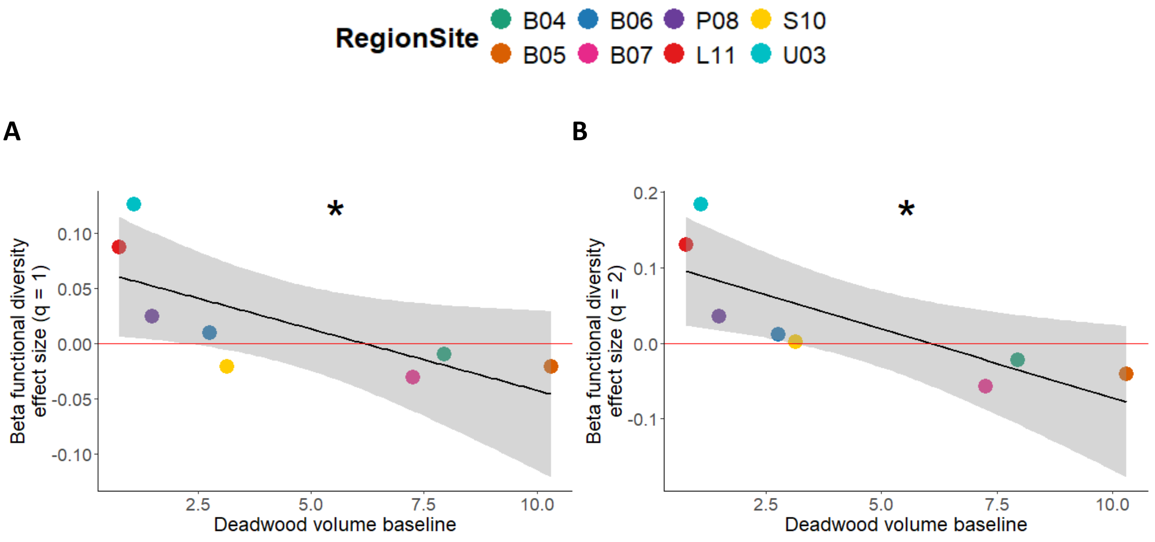


Figure S16: Regression plots of significant ($p < 0.05$) linear relationships between diversity measure effect sizes due to ESBC and deadwood volume. (A) β functional diversity, q = 1; (B) β functional diversity, q = 2.

Table 2S: Weights assigned to each forest site in the robust regression models across all significant response–predictor combinations. Each cell shows the weight assigned to a given site for a specific model. Sites with weights below 0.8 are considered potential outliers for that model.

| **response** | **predictor** | **B04** | **B05** | **B06** | **B07** | **L11** | **P08** | **S10** | **U03** |
|---|---|---|---|---|---|---|---|---|---|
| Gamma TD effect size (q = 0) | Soil pH baseline | 0.74 | 0.48 | 1.00 | 1.00 | 1.00 | 1.00 | 1.00 | 1.00 |
| Gamma TD effect size (q = 0) | Understory biomass baseline | 1.00 | 0.68 | 1.00 | 1.00 | 1.00 | 1.00 | 1.00 | 1.00 |
| Gamma TD effect size (q = 0) | Understory species richness baseline | 0.45 | 0.21 | 0.73 | 1.00 | 1.00 | 1.00 | 1.00 | 1.00 |
| Gamma TD effect size (q = 0) | Soil C:N ratio baseline | 1.00 | 0.30 | 1.00 | 1.00 | 1.00 | 0.54 | 1.00 | 1.00 |
| Beta TD effect size (q = 1) | Understory biomass baseline | 1.00 | 0.58 | 1.00 | 1.00 | 1.00 | 1.00 | 0.29 | 1.00 |
| Beta TD effect size (q = 2) | Understory biomass difference | 1.00 | 1.00 | 1.00 | 1.00 | 0.23 | 1.00 | 1.00 | 1.00 |
| Beta TD effect size (q = 2) | Soil pH baseline | 1.00 | 1.00 | 1.00 | 1.00 | 0.59 | 1.00 | 0.49 | 1.00 |

| | | | | | | | | | |
|---|---|---|---|---|---|---|---|---|---|
| Beta TD effect size (q = 2) | Understory biomass baseline | 0.95 | 1.00 | 0.80 | 1.00 | 1.00 | 1.00 | 0.99 | 1.00 |
| Beta TD effect size (q = 2) | Understory species richness baseline | 1.00 | 1.00 | 1.00 | 1.00 | 0.48 | 1.00 | 1.00 | 1.00 |
| Beta TD effect size (q = 2) | Soil sand fraction baseline | 1.00 | 1.00 | 1.00 | 1.00 | 1.00 | 1.00 | 1.00 | 1.00 |
| Gamma FD effect size (q = 1) | Soil C:N ratio baseline | 1.00 | 1.00 | 1.00 | 1.00 | 1.00 | 1.00 | 1.00 | 1.00 |
| Gamma FD effect size (q = 2) | Soil C:N ratio baseline | 1.00 | 1.00 | 1.00 | 1.00 | 1.00 | 1.00 | 1.00 | 1.00 |
| Alpha FD effect size (q = 1) | Soil pH baseline | 1.00 | 1.00 | 1.00 | 1.00 | 1.00 | 1.00 | 0.74 | 1.00 |
| Alpha FD effect size (q = 1) | Understory biomass baseline | 1.00 | 1.00 | 1.00 | 1.00 | 1.00 | 1.00 | 1.00 | 1.00 |
| Alpha FD effect size (q = 1) | Understory species richness baseline | 1.00 | 1.00 | 1.00 | 1.00 | 1.00 | 1.00 | 1.00 | 1.00 |
| Alpha FD effect size (q = 1) | Soil C:N ratio baseline | 1.00 | 1.00 | 1.00 | 1.00 | 1.00 | 1.00 | 1.00 | 1.00 |
| Alpha FD effect size (q = 1) | Soil sand fraction baseline | 1.00 | 1.00 | 1.00 | 1.00 | 1.00 | 1.00 | 1.00 | 1.00 |
| Alpha FD effect size (q = 2) | Soil pH baseline | 1.00 | 1.00 | 1.00 | 1.00 | 0.69 | 1.00 | 1.00 | 1.00 |
| Alpha FD effect size (q = 2) | Understory biomass baseline | 1.00 | 1.00 | 1.00 | 1.00 | 1.00 | 1.00 | 1.00 | 1.00 |
| Alpha FD effect size (q = 2) | Understory species richness baseline | 1.00 | 1.00 | 1.00 | 1.00 | 0.74 | 1.00 | 1.00 | 1.00 |
| Alpha FD effect size (q = 2) | Soil C:N ratio baseline | 1.00 | 1.00 | 1.00 | 1.00 | 0.89 | 1.00 | 1.00 | 1.00 |
| Alpha FD effect size (q = 2) | Soil sand fraction baseline | 1.00 | 1.00 | 1.00 | 1.00 | 1.00 | 1.00 | 1.00 | 1.00 |
| Beta FD effect size (q = 1) | Deadwood volume baseline | 1.00 | 1.00 | 1.00 | 1.00 | 1.00 | 1.00 | 0.91 | 0.62 |
| Beta FD effect size (q = 1) | Understory biomass baseline | 1.00 | 1.00 | 1.00 | 1.00 | 1.00 | 1.00 | 1.00 | 1.00 |
| Beta FD effect size (q = 1) | Understory species richness baseline | 1.00 | 1.00 | 1.00 | 1.00 | 0.55 | 1.00 | 1.00 | 1.00 |

| | | | | | | | | | |
|---|---|---|---|---|---|---|---|---|---|
| Beta FD effect size (q = 1) | Soil sand fraction baseline | 1.00 | 1.00 | 1.00 | 1.00 | 1.00 | 0.78 | 0.92 | 1.00 |
| Beta FD effect size (q = 2) | Deadwood volume baseline | 1.00 | 1.00 | 1.00 | 1.00 | 1.00 | 1.00 | 1.00 | 0.83 |
| Beta FD effect size (q = 2) | Understory biomass baseline | 1.00 | 1.00 | 1.00 | 1.00 | 1.00 | 1.00 | 1.00 | 1.00 |
| Beta FD effect size (q = 2) | Understory species richness baseline | 1.00 | 1.00 | 1.00 | 1.00 | 0.52 | 1.00 | 1.00 | 1.00 |
| Beta FD effect size (q = 2) | Soil sand fraction baseline | 1.00 | 1.00 | 1.00 | 0.83 | 1.00 | 0.56 | 1.00 | 1.00 |
